# OEF18 is a membrane-anchored organellar Ca²⁺ sensor linking calcium signaling to jasmonate-mediated defense and stress acclimation

**DOI:** 10.64898/2026.08.13.744648

**Authors:** Simon Stael, Przemyslaw Kmiecik, Bernhard Wurzinger, Shuning Qi, Dominic Kuang, Elena Sanchez Martin-Fontecha, Roman Bayer, Barbara Pfister, Michael Reichelt, Ingo Ebensberger, Inge De Clercq, Axel Mithöfer, Markus Teige

## Abstract

Changes in intracellular calcium ion (Ca²⁺) concentrations generate characteristic signatures that are decoded by specialized Ca²⁺-binding proteins (CaBP). Although substantial progress has been made in understanding cytosolic calcium signaling pathways, calcium signaling within organelles, particularly chloroplasts, remains poorly understood, partly because only a few EF-hand CaBP have been identified in organelles. Here, we describe a novel EF-hand protein of 18 kDa, that was found to be associated with the chloroplast envelope and peroxisomal membrane and was therefore named OEF18 (ORGANELLAR EF-HAND PROTEIN OF 18 kDa). OEF18 has a very unusual structure, containing an N-terminal myristoylation site, followed by one EF-hand in the N-terminus facing to the cytosol, and a transmembrane domain in the C-terminus. OEF18 membrane-targeting was found to be mediated by ANKYRIN REPEAT-CONTAINING PROTEIN 2A (AKR2A) via the C-terminal transmembrane domain of OEF18. Furthermore, the EF-hand in OEF18 bound Ca²⁺ at a physiological concentration that led to a large protein conformational change, inducing oligomerization of the N-terminal part. We found that *oef18* mutants accumulated less jasmonic acid (JA) and its bioactive conjugate JA-Ile, likely causing a defect in the insect herbivore response. Wild-type OEF18 complemented the herbivory phenotype of *oef18* mutants, whereas an EF-hand point mutant lacking Ca²⁺-binding capacity failed to restore the wild-type response. Furthermore, OEF18 was required for resistance to salt stress in combination with dark-induced senescence. Together, these results establish OEF18 as a previously unrecognized organellar Ca²⁺ sensor that couples Ca²⁺ perception to JA-mediated defense and abiotic stress responses in plants.

## Introduction

Ca^2+^ is a universal second messenger acting in the regulation of growth and developmental processes as well as responses to environmental conditions (Luan, 2026; Luo *et al*., 2026). At mM levels, Ca^2+^ can precipitate with phosphate (Pi), thereby disrupting energy metabolism and many other cellular processes (Carafoli and Krebs, 2016). Hence, all living cells keep the cytosolic Ca^2+^ concentration low, typically in the range of 50-100 nM. This is achieved by active transport of Ca^2+^ to stores, including the vacuole, the endoplasmic reticulum (ER) and the apoplast (Z., Li *et al*., 2023; Luan and Wang, 2021). As a consequence, the large gradient of Ca^2+^ between the stores and the cytosol allows for a fast passive release followed by active reuptake, leading to the generation of so called Ca^2+^ signals or signatures (Berridge *et al*., 2000; Luan, 2026; Clapham, 2007).

Ca²⁺-binding proteins (CaBPs) are at the center of decoding these signals, often inducing conformational changes or altering protein-protein interactions, thereby initiating downstream signaling events. In plants, one of the most prominent groups of CaBPs comprises the EF-hand proteins (Luan and Wang, 2021). The EF-hand is a conserved helix-loop-helix structural motif that coordinates Ca²⁺ through specific amino acid residues within the loop region and typically binds Ca²⁺ with low capacity but high affinity. They can be divided into calmodulins (CaMs), calmodulin-like proteins (CMLs), calcineurin B-like proteins (CBLs) and as decoders of calcium signals, calcium-dependent protein kinases (CDPKs/CPKs), CPK-related protein kinases (CRKs), and calcium- and calmodulin-dependent protein kinases (CCaMKs) (Luan, 2026). Recently completed large sequencing projects revealed that genomes of different plant and algal species encode on average about 1-5 CaMs, 10-30 CMLs, 2-14 CBLs, and 2-53 CPKs (Li *et al*., 2023; Mohanta *et al*., 2019).

Early transcriptomics studies revealed that many of the genes encoding these EF-hand proteins are stress-responsive, suggesting a role in Ca²⁺-mediated stress signaling (McCormack *et al*., 2005; Zhu *et al*., 2015). For example, in the salt oversensitive (SOS) pathway, the response to salt stress is mediated by CBLs as Ca²⁺ sensors that interact with protein kinases (CIPKs) and activate Na^+^ transporters such as SOS1 or NHX1 (Qiu *et al*., 2002; Steinhorst *et al*., 2022). Similarly, the CPK family was implicated in the salt stress response (Mehlmer *et al*., 2010). Furthermore, the CML family was shown to play important roles during herbivore resistance, innate immunity or salt stress, or during both abiotic and biotic stresses (Zhu *et al*., 2015). Ca²⁺ is also heavily involved in antiviral defense (Zvereva *et al*., 2024) and pathogens exploit this by targeting calcium sensors such as CaM and CMLs with their secreted effectors (Galaud *et al*., 2025), often specifically affecting chloroplasts (Zvereva *et al*., 2024). While research traditionally focused on cytosolic Ca²⁺ signaling cascades, increasing evidence indicates that organelles generate and decode distinct Ca²⁺ signals (Stael *et al*., 2012; Luan, 2026; Pirayesh *et al*., 2021). Among these organelles, chloroplasts have emerged as important hubs of Ca²⁺ signaling (Corti *et al*., 2023; Navazio *et al*., 2020), integrating environmental cues such as light-dark transitions (Frank *et al*., 2019; Martí Ruiz *et al*., 2020; Sai and Johnson, 2002), high-light exposure (Kuang *et al*., 2025; Flori *et al*., 2025; Pivato *et al*., 2023), heat shock (Lenzoni and Knight, 2019; Kuang *et al*., 2025), and pathogen infection (Nomura *et al*., 2012). Ca^2+^ was also shown to regulate chloroplast protein import (Chigri *et al*., 2005) via CaM-binding to the inner envelope translocon component Tic32 (Chigri *et al*., 2006). At the molecular level, the Calcium sensing receptor (CAS) plays a central role (Li *et al*., 2022). CAS was initially characterized as a cell-surface Ca^2+^ receptor that, unlike EF-hands, binds Ca²⁺ with low affinity but high capacity (Han *et al*., 2003). Subsequent chloroplast proteomic analyses and functional studies revealed CAS as a thylakoid-localized protein required for Ca^2+^-dependent stomatal responses whose phosphorylation state is directly regulated by light intensity (Vainonen *et al*., 2008; Friso *et al*., 2004; Peltier *et al*., 2004; Weinl *et al*., 2008; Nomura *et al*., 2008). Upon biotic challenge, a CAS-dependent chloroplastic Ca²⁺ transient acts as an upstream trigger driving plant immune signaling and salicylic acid-dependent defenses (Nomura *et al*., 2012). This highlights the critical need to understand how Ca²⁺ signals are generated, perceived and transmitted within chloroplasts.

Compared to the cytosol, CaBPs in chloroplasts are underrepresented. Based on ∼250 predicted EF-hand proteins in the *A. thaliana* proteome and assuming a uniform subcellular distribution (chloroplast proteome representing ∼9 % of the total proteome), one would expect ∼23 EF-hand proteins to localize to chloroplasts. Yet only a single EF-hand protein has so far been experimentally detected within the chloroplast stroma: the Ca²⁺-activated RelA/SpoT homolog protein (CRSH), which is unique in combining two EF-hand motifs with a RelA/SpoT enzymatic domain. Upon Ca²⁺ binding, CRSH regulates the synthesis of the alarmone guanosine 5′-diphosphate 3′-diphosphate (ppGpp) (Romand *et al*., 2022; Masuda *et al.,* 2008).

Searching for novel chloroplast-localized CaBPs, we previously described the localization of CALCIUM-DEPENDENT PROTEIN KINASE16 (CPK16) (Stael, Bayer, *et al*., 2011). Although CPK16 contains a predicted chloroplast transit peptide, it exhibits dual localization to either the plasma membrane or the chloroplast stroma, depending on the presence or absence of N-terminal myristoylation, respectively. Myristoylation is the covalent attachment of a C14 fatty acid myristate to an N-terminal glycine residue and serves as an important mechanism for membrane association and subcellular targeting. Notably, this lipid modification can be exploited by viral pathogens to promote membrane recruitment, intracellular trafficking, and infection, including in CPK16 (Medina-Puche *et al*., 2020). To identify additional chloroplast proteins that may undergo myristoylation, N-myristoylation probabilities were predicted for proteins with experimentally supported chloroplast localization based on mass spectrometry data in the Plant Proteome Database (Sun *et al*., 2009; Bologna *et al*., 2004). During this search, we came across a protein of unknown function (At1g64850) predicted to have an EF-hand, which we named ORGANELLAR EF-HAND PROTEIN OF 18 kDa (OEF18).

## Results

### OEF18 predominantly localizes to the chloroplast envelope and is potentially dually localized at the peroxisomal membrane

In an early study, the rice ortholog of OEF18 was first identified by proteomics in the plastid envelope of etiolated rice seedlings (Zychlinski *et al*., 2005). Since then, *A. thaliana* OEF18 was observed in three additional proteomic studies of the plastid envelope (Bräutigam and Weber, 2009; Bouchnak *et al*., 2019; Ferro *et al*., 2010). Furthermore, OEF18 was identified by proteomics in the mitochondria (Fuchs *et al*., 2020) and peroxisomes (Reumann *et al*., 2009). In accordance with these studies, an OEF18-YFP fusion protein co-localized with the chloroplast envelope protein OUTER ENVELOPE MEMBRANE PROTEIN 7 (OEP7) in transient tobacco (*Nicotiana tabacum*) expression assays (Fig. 1A). Additional undefined accumulations of OEF18 were found outside of chloroplasts. Transient co-expression of OEF18-YFP with fluorescent organellar marker proteins confirmed the presence of OEF18 in peroxisomes (Fig. 1B), particularly in the peroxisome membrane, judged by the ring-like accumulations of OEF18-YFP at peroxisomes (Fig. 1C). No conclusive evidence could be obtained for co-localization with mitochondria (Fig. S1A).

**Figure 1.**
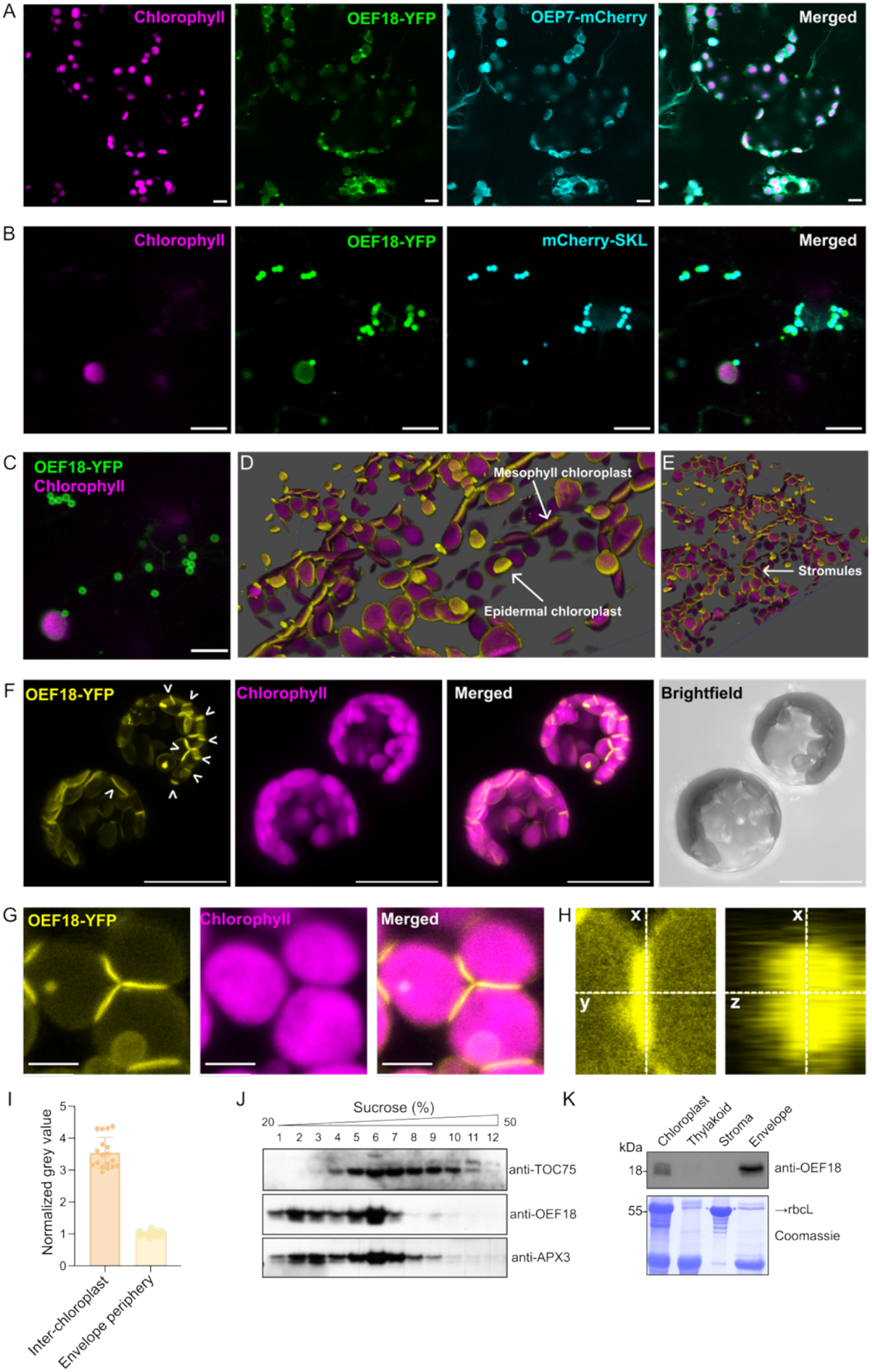
OEF18 displays chloroplast envelope and peroxisomal membrane localization. (A, B) Confocal microscopy images of colocalization of OEF18-YFP and the plastid outer envelope marker OEP7-mCherry (A) or the peroxisome marker mCherry-SKL (B) in *N. tabacum* epidermal cells. Scale bar, 20 μm. (C) Image of the sample in B at lower laser excitation reveals ring-like accumulation of OEF18-YFP surrounding the peroxisome. Scale bar, 20 μm. (D, E) Three-dimensional (3D) reconstructions of OEF18-YFP localization (yellow) in *A. thaliana* epidermal and mesophyll cells. Chlorophyll autofluorescence in magenta. (F) Z projection of an OEF18-YFP image stack from *A. thaliana* protoplasts. White arrowheads indicate inter-chloroplast OEF18-YFP signal accumulation (top right protoplast), or localization of OEF18-YFP to a narrow band around the equator of the chloroplast. Scale bar, 20 μm. (G, H) Higher-magnification images and orthogonal views (x-y and x-z) of OEF18-YFP accumulation relative to chloroplasts. Scale bar, 5 μm. (I) Signal intensity (normalized gray values) of OEF18-YFP in the inter-chloroplast and envelope periphery. (J) Western blot of sucrose density gradient fractions (20%-50%) indicating the membrane association of OEF18 in *A. thaliana* leaves relative to a panel of organelle marker proteins. TOC75 from the top panel and APX3 from the bottom panel were used as markers for chloroplast outer envelope membrane and peroxisomal membrane, respectively. (K) Western blot of intact *A. thaliana* chloroplasts and thylakoid, stroma, and envelope fractions. Coomassie stain of the rubisco large subunit (rbcL) indicates the fractionation worked.

Two *A. thaliana* T-DNA insertion mutant lines, WiscDsLox461-464A5 and SK40148, were identified and verified to be loss-of-function mutants by PCR and western blot analysis with an OEF18-specific antibody, and named *oef18-1* and *oef18-2*, respectively (Fig. S2A-D). Expression of OEF18-YFP driven by an upstream 1 kb promoter fragment in the *oef18-1* background (pOEF18::OEF18-YFP/*oef18-1*, abbreviated as OEF18-YFP^c^) led to a similar localization at the chloroplast envelope. In these stable *A. thaliana* lines, the OEF18-YFP fusion protein preferentially accumulates at the equator of chloroplasts, forming a prominent ’belt’ around the flattened organelle (Fig. 1D-F). This belt was most obvious in mesophyll cells, but a similar localization pattern was also observed in epidermal chloroplasts. Furthermore, OEF18-YFP occasionally localized to distinct disc-like accumulations at the interface of chloroplasts in protoplasts (Fig. 1F-H) and intact leaf cells (Fig. S1B and C). Theoretically, a mere overlap of two adjacent envelopes would lead to a twofold YFP signal intensity. However, OEF18-YFP appears enriched at these interfaces, as YFP intensity is higher than threefold compared to adjacent regions of the same chloroplasts (Fig. 1I). As peroxisomes can adopt elongated shapes (Reumann and Bartel, 2016), we initially hypothesized that these inter-chloroplast OEF18-YFP accumulations might represent peroxisomes compressed between chloroplasts. To verify, we stably transformed the OEF18-YFP^c^ line with a peroxisomal marker (mScarlet-SKL). Surprisingly, the OEF18-YFP inter-chloroplast accumulations did not co-localize with peroxisomes. In contrast to transient expression in tobacco, we did not observe any co-localization of OEF18-YFP with peroxisomes in stable *A. thaliana* lines (Fig. S1B and C).

To complement the microscopy localization data, a biochemical fractionation of leaf protein extracts was performed by sucrose gradient centrifugation followed by western blot against OEF18 and subcellular membrane markers (Fig. 1J). From the twelve fractions, OEF18 seemed to overlap best with ASCORBATE PEROXIDASE 3 (APX3), which localizes to the peroxisomal membrane (Shen *et al*., 2010), and to a lesser extent with TRANSLOCON AT THE OUTER ENVELOPE MEMBRANE OF CHLOROPLASTS 75 (TOC75). Fractionation of isolated chloroplasts from wild type *A. thaliana* plants indicated that endogenous OEF18 protein accumulates in the envelope fraction (Fig. 1K). Taken together, OEF18 predominantly localizes to the plastid envelope, while evidence from transient expression and fractionation experiments suggests a possible, but not yet fully resolved, association with peroxisomal membranes.

### The N-terminal EF-hand and C-terminus of OEF18 face the cytosol

OEF18 has two predicted transmembrane helices according to the ARAMEMNON 8.1 database (Schwacke *et al*., 2003) with the protein N-terminus containing the predicted EF-hand domain (Fig. 2A). To ascertain the membrane topology of OEF18, we used an assay based on co-expression of self-assembly GFP in transient tobacco assays (Wiesemann *et al*., 2013). This assay is conceptually similar to the well-established bimolecular fluorescence complementation (BiFC) approach. However, GFP is divided into two unequal fragments, β-strands 1-10 (GFP1-10) and strand 11 (GFP11), which exhibit higher intrinsic affinity than BiFC fragments. This high affinity enables GFP reconstitution when both fragments are present in the same cellular compartment, independent of any interaction between the fused proteins. Comparison with proteins of known membrane topology or subcellular localization allows the topology of the protein of interest to be inferred. OEP7 has a single transmembrane helix with the C-terminus located in the cytosol (Schleiff *et al*., 2001; Sommer *et al*., 2011). Fusion of GFP11 to either the OEF18 N-terminus or C-terminus resulted in the interaction with a GFP1-10 fusion protein of OEP7 at the chloroplast envelope when fused to the C-terminus (Fig. 2B). The N- and C-terminus of OEF18 are likely situated at the same side of the membrane as an N- and C-terminal fusion of OEF18 with GFP11 and GFP1-10, respectively, resulted in reconstituted GFP signals at the chloroplast envelope and accumulations that are presumably peroxisomes (Fig. 2C). Furthermore, GFP1-10 fusion proteins with cytosolic proteins CASEIN KINASE 1A (CK1A) and NITRATE REDUCTASE 2 (NIA2) interacted with N- and C-terminal GFP11 fusion proteins of OEF18 at the chloroplast envelope and peroxisome-like accumulations (Fig. S3). Together this indicates that both the N-terminus, containing the EF-hand, and C-terminus of OEF18 are localized in the cytosol.

**Figure 2.**
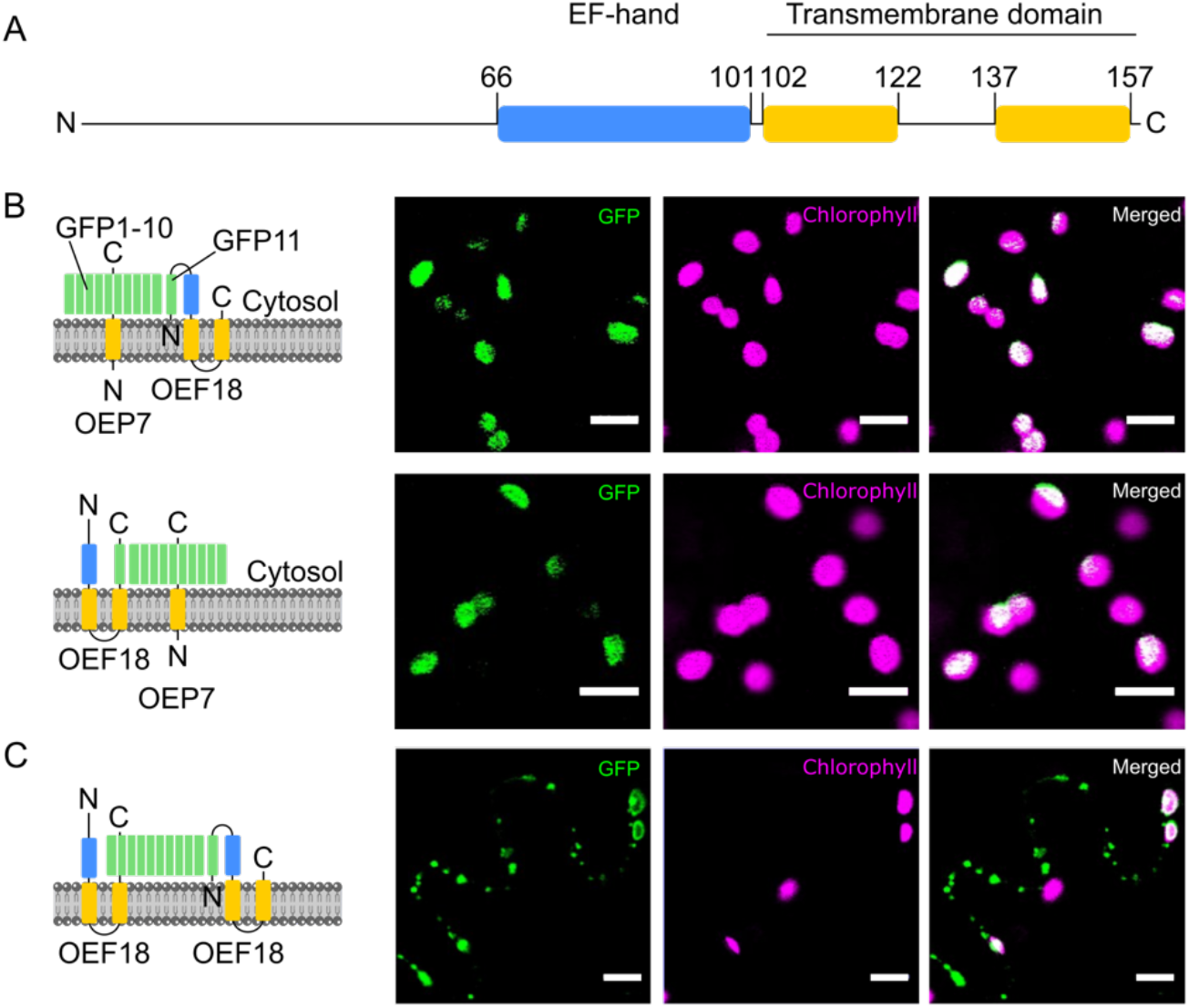
Both the N-terminus, containing the EF-hand, and C-terminus of OEF18 are localized in the cytosol. (A) Position of the predicted EF-hand and transmembrane domain in the OEF18 amino acid sequence. (B) Topology assessment of OEF18 by saGFP. Confocal microscopy images of *N. tabacum* cells co-expressing either the chloroplast outer envelope proteins OEP7_1-10C_ together with OEF18_11N_ (lines 1) or OEF18_11c_ (lines 2). (C) *N. tabacum* cells co-expressing OEF18_1-10C_ and OEF18_11N_. Scale bar, 20 μm.

### AKR2A interaction and N-myristoylation are involved in the membrane targeting of OEF18

OEF18 lacks a strong canonical targeting signal for either chloroplasts or peroxisomes. However, tail-anchored (TA) proteins are known to be targeted to both chloroplast envelope and peroxisomal membranes in the absence of canonical chloroplast- or peroxisome-targeting signals. Although OEF18 contains two predicted transmembrane helices, the presence of positively charged residues C-terminal to the first transmembrane helix, including a Lys-Lys motif in position 125-126 and Lys-136, is reminiscent of sequence features typical of TA proteins (Lee *et al*., 2011). Chloroplast envelope membrane targeting of such proteins may be mediated by AKR2A, which recognizes hydrophobic transmembrane domains together with the C-terminal positively charged region (CPR) in TA proteins (Bae *et al*., 2008; Shen *et al*., 2010). AKR2A binds protein clients co-translationally (Kim *et al*., 2015) and, in conjunction with additional cofactors such as Hsp17.8 (Kim *et al*., 2011), facilitates their delivery to specific membrane systems. We tested a potential interaction between AKR2A and OEF18 using a yeast two-hybrid assay (Fig. 3A and 3B). OEP7 and PMP22 were used as positive and negative controls, respectively, however, the interaction pattern differed partially from previous reports (Bae *et al*., 2008; Shen *et al*., 2010). When AKR2A was fused to the LexA DNA-binding domain, PMP22 showed a detectable interaction with AKR2A, whereas OEP7 did not exhibit interaction above the empty-vector control (Fig. 3A). In contrast, when AKR2A was fused to the Gal4 activation domain, OEP7 interacted more strongly with AKR2A than PMP22, consistent with earlier studies (Fig. 3B). Despite this orientation-dependent behavior of the control proteins, full-length OEF18 interacted with AKR2A in both bait/prey configurations. Importantly, an N-terminal OEF18 fragment lacking the transmembrane domain failed to interact with AKR2A, indicating that the C-terminal membrane-anchoring region is required for the interaction (Fig. 3A and 3B).

**Figure 3.**
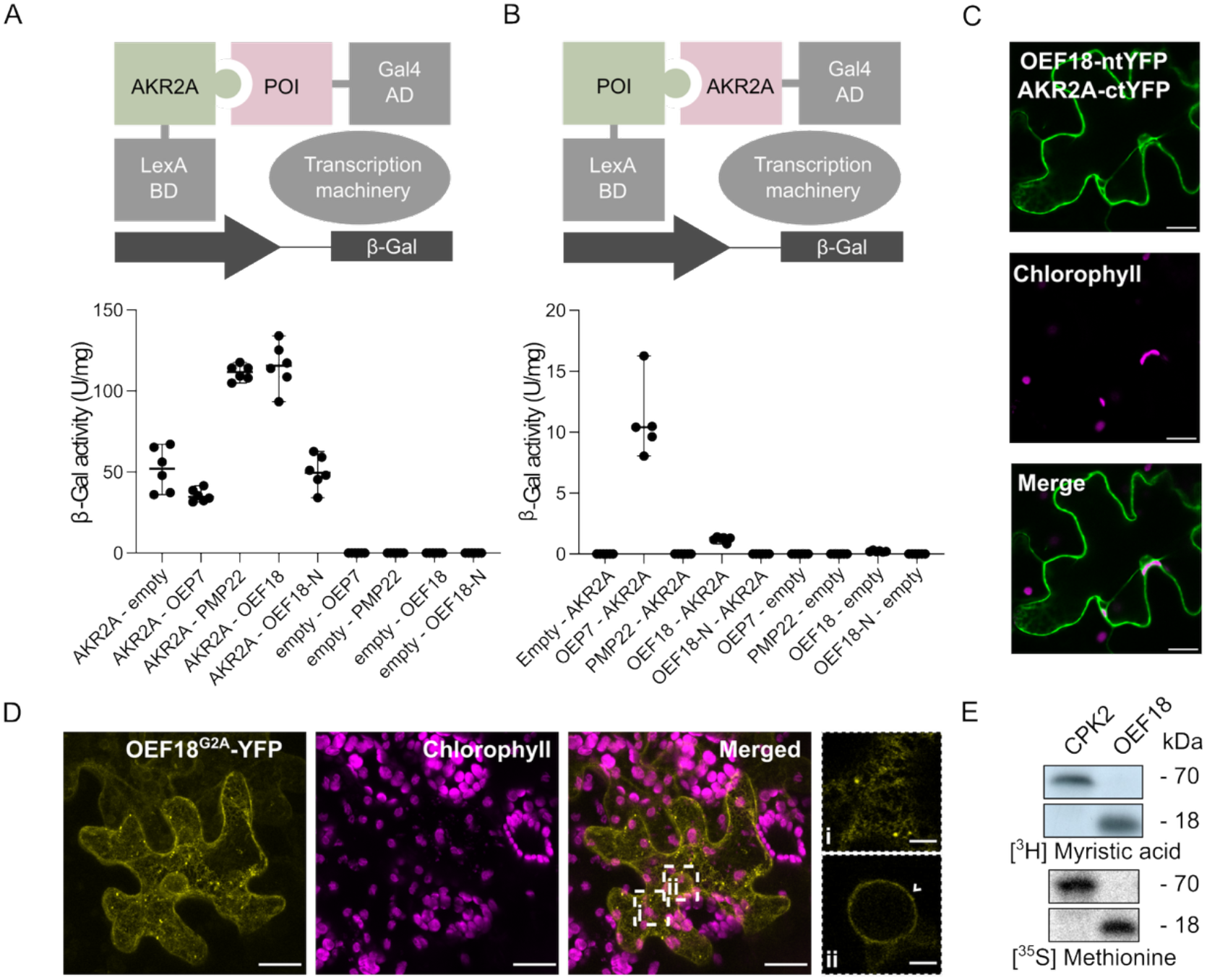
OEF18 interacts with AKR2A and undergoes N-terminal myristoylation required for proper subcellular localization. (A, B) A schematic diagram of the yeast two-hybrid system, in which AKR2A or the protein of interest (POI) were fused to the GAL4 activation domain (AD) or the LexA binding domain (BD) to test protein-protein interaction. OEP7 and PMP22 were used as controls. (C) Confocal microscopy image of OEF18 fused to the N-terminal fragment of YFP (nYFP) and AKR2A fused to the C-terminal fragment of YFP (cYFP) co-expressed in tobacco leaf epidermis cells. Scale bar, 20 μm. (D) Confocal image of OEF18-YFP in which the N-terminal Gly is mutated to Ala (OEF18^G2A^) in tobacco leaf epidermis cells. Scale bar, 20 μm. (E) Radiolabeled myristoylation assay of CPK2 (positive control) and OEF18 in wheat germ lysate.

These findings were further validated by BiFC assays between AKR2A and OEF18 in tobacco transient expression assays (Fig. 3C). Consistent with a co-translational interaction and binding of AKR2A to the transmembrane helix of client proteins, reconstituted YFP signal of the AKR2A-OEF18 interaction was found in the cytosol. Furthermore, transiently expressed OEF18 carrying mutations in the Lys-Lys motif (substituted with Ile and Glu; OEF18^KK2IE^) was partially mistargeted to a compartment resembling the ER (Fig. S4C). Similar mis-targeting to the ER was reported previously when mutating the CPR (Lee *et al*., 2011).

OEF18 is predicted to be N-myristoylated (Bologna *et al*., 2004) and was identified in the *A. thaliana* myristoylated proteome (Majeran *et al*., 2018). Substitution of Gly-2, the predicted N-myristoylation site, with Ala (OEF18^G2A^) caused severe mistargeting and a more diffuse subcellular distribution of OEF18 (Fig. 3D), resembling an ER-like localization similar to the OEF18^KK2IE^ mutant. Myristoylation of OEF18 was confirmed *in vitro* by expression of full-length OEF18 in wheat germ lysate in the presence of [³H]-labeled myristate. CALCIUM-DEPENDENT PROTEIN KINASE 2 (CPK2) served as a positive control for myristoylation (Lu and Hrabak, 2002) and incorporation of [^35^S]-methionine served as protein expression control (Fig. 3E). Evolutionary conservation of the Lys-Lys motif and Gly-2 reinforce the importance of these amino acids for the correct targeting of OEF18 (Fig. S5). In several orthologs, however, we find that the Lys is substituted by an Arg, indicating a requirement for a positively charged amino acid rather than strictly Lys. Together, these results indicate that OEF18 targeting depends on AKR2A-mediated recognition and requires both an intact CPR and N-terminal myristoylation for correct membrane localization.

TA proteins are generally not myristoylated. A comparison of TA proteins from Brito *et al*., (2019) and myristoylated proteins from Majeran *et al*. (2018) revealed only four overlapping proteins, including two disease resistance proteins and two proteins of unknown function (Fig. S6A). Furthermore, only few proteins including CALCIUM-DEPENDENT PROTEIN KINASE 1 (CPK1) and ANTHER DEHISCENCE REPRESSOR (ADR) are known to be both myristoylated and to localize to peroxisomes (Fig. S6B) (Dai *et al*., 2019). Similarly, only few chloroplast-localized proteins are myristoylated (Fig. S6C). Relative to myristoylation, TA proteins were present more frequently in chloroplasts and peroxisomes (Fig. S6D and E; Kriechbaumer *et al*., 2009). Altogether, the combination of TA protein characteristics and myristoylation in OEF18 is an unusual but important feature for correct localization, as Gly-2 and the transmembrane domain are deeply conserved (Fig. S5).

### OEF18 is ubiquitously expressed, evolutionarily conserved, and has a paralog, OEF18L, that lacks a functional EF-hand

According to the *A. thaliana* eFP browser (Sullivan *et al*., 2019), *OEF18* is expressed in most plant organs and developmental stages (Fig. 4A). To validate the distribution of OEF18 protein in *A. thaliana*, we performed immunoblot analyses using an anti-OEF18 antibody, whose specificity was verified in the *oef18* knockouts and complementation lines (Fig. S2D and E). Tissue-specific analysis revealed that OEF18 is present in major plant organs, but is absent from seeds (Fig. 4B). Furthermore, OEF18 accumulation appears to be associated with leaf development, as assessed across leaves of different ages. A progressive increase in OEF18 protein levels was observed from young (leaf 6) to old (leaf 1) rosette leaves (Fig. 4C). This gradient suggested that *OEF18* expression is developmentally regulated and may play a more prominent role during late stages of leaf development. Indeed, OEF18 levels were highest in old and senescent leaves compared to cotyledons, young rosette leaves (2-week-old), and old rosette leaves (4-week-old; Fig. 4D). These results suggest that OEF18 is present in both vegetative and reproductive tissues but can be enriched in specific cell types or developmental contexts, hinting at a potential role in senescence.

**Figure 4.**
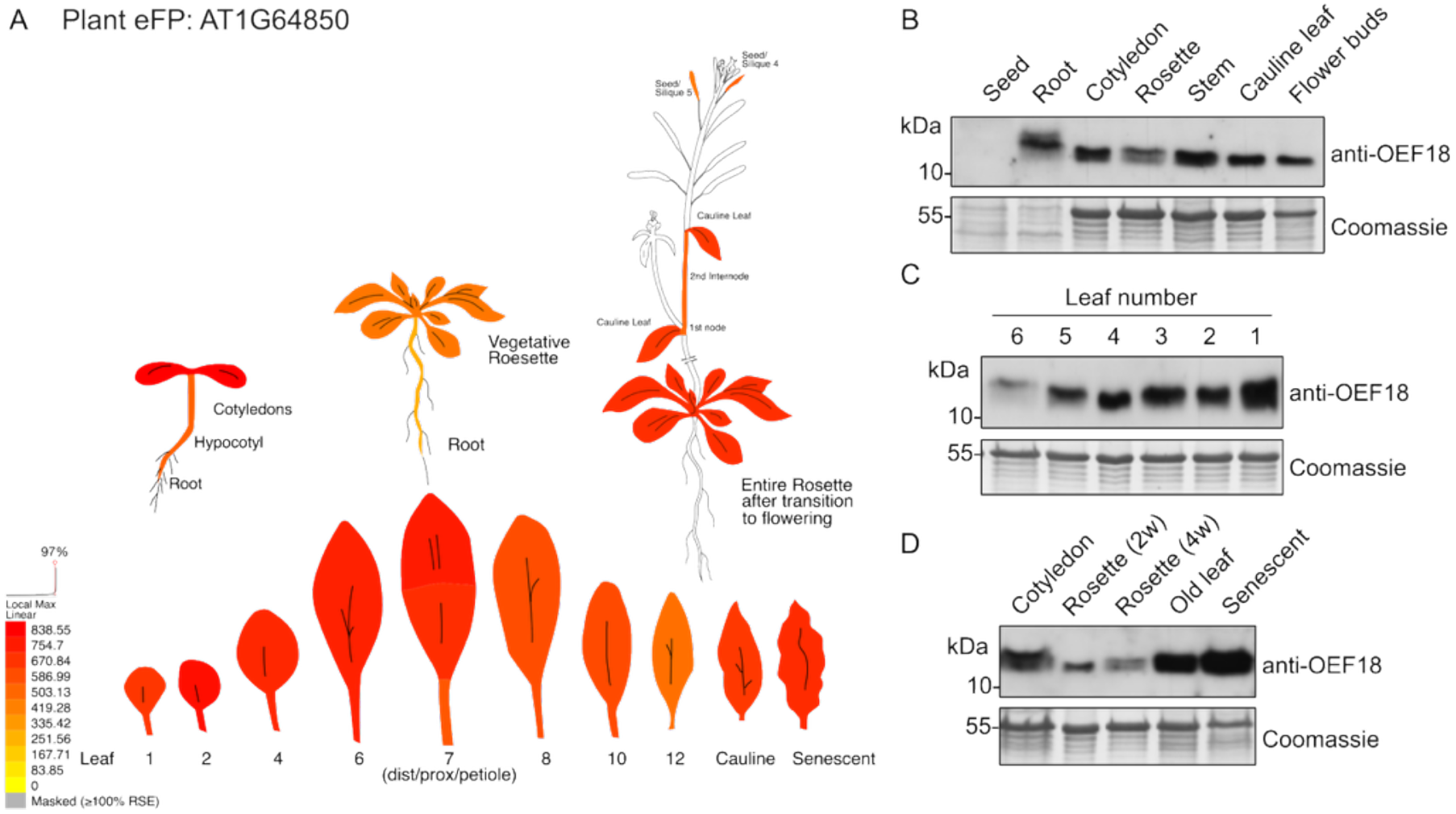
OEF18 is ubiquitously expressed in various *A. thaliana* tissues. (A) Expression pattern of OEF18 across *A. thaliana* tissues based on ePlant database visualization. (B-D) Western blot against OEF18 in *A. thaliana* of various plant organs (B), rosette leaves of increasing developmental stages (C), and leaves of different developmental and senescence stages, including cotyledons (10 days), 2-week-old and 4-week-old rosette leaves, old leaves (2 months) and senescent leaves (clearly yellowing) (D). Protein load is indicated by Coomassie Brilliant Blue stain.

We next investigated the taxonomic distribution of OEF18 and its evolutionary trajectory. To this end, we first used a BlastP search in the proteome of *A. thaliana* to screen for the existence of paralogs to this protein. This revealed a single protein, AT4G37445, that retained the TM1-KR-TM2 domain architecture but lacks several key Ca²⁺-coordinating residues at the site corresponding to the EF-hand motif in OEF18 (Fig. S7A). Similar to OEF18, this protein localized to the chloroplast envelope and co-localized with a peroxisome marker in transient expression assays in tobacco (Fig. S7B). Therefore, we named this protein OEF18-LIKE (OEF18L).

Subsequently, we performed a targeted search for OEF18 and OEF18L orthologs (Table S1). OEF18 orthologs were identified across the *Streptophyta* and also within the *Chlorophyta*, and retained the conserved protein architecture, including the EF-hand domain and the adjacent transmembrane helices (TM1 and TM2) flanking the KK/R motif (Fig. S5). Orthologs to OEF18L were confined to the flowering plants. We computed the combined maximum likelihood (ML) tree with OEF18 and OEF18L orthologs (Fig. 5A). Consistent with the detection of OEF18L orthologs only in the flowering plants, these sequences formed a monophyletic clade in the tree. However, the precise timing of the gene duplication event giving rise to the ortholog pair OEF18 - OEF18L remained unclear. The OEF18 orthologs of *Chara braunii* grouped basal to the OEF18L clade to the exclusion of OEF18 orthologs in the flowering plants. Moreover, the OEF18 orthologs of *Physcomitrium patens* and *Selaginella moellendorffii* formed a clade together with the OEF18 orthologs of the flowering plants and to the exclusion of OEF18L. If taken at face value, both findings would suggest that the gene duplication occurred prior to the diversification of the *Streptophyta* and was followed by mutual gene loss in the basal *Streptophyta* lineages. However, bootstrap values of the corresponding splits are low, indicating that the phylogenetic signal in the data is low. We therefore tested whether the ML tree explained the data significantly better than a tree that placed the orthologs of *S. moellendorffii*, *P. patens* and *C. braunii* at the base of the *Streptophyta* and prior to the emergence of the two paralogous groups. The ML tree failed to reject this alternative hypothesis (AU-test: p > 0.05) (Fig. 5B). Hence both the phylogenetic profiles of OEF18 and OEF18L and the phylogenetic signal in the respective orthologous sequences was consistent with a gene duplication event on the lineage of the flowering plants after the diversification of *P. patens* and *S. moellendorffii*. These data collectively suggest that OEF18 is evolutionarily ancient, while OEF18L is a more recently evolved paralog that likely lost its Ca^2+^-binding capacity but preserved its original subcellular localization.

**Figure 5.**
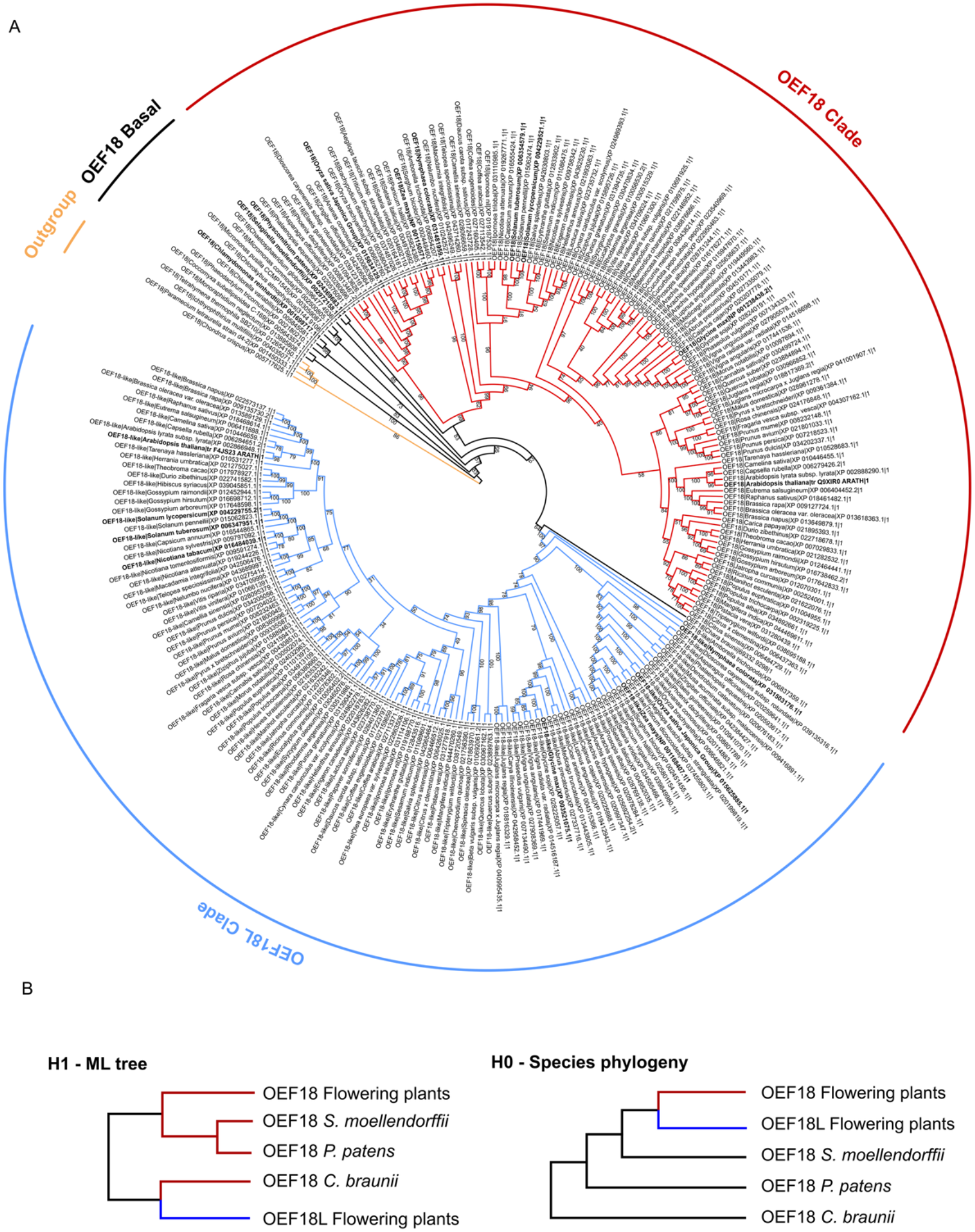
Maximum likelihood phylogenetic tree reveals the evolutionary divergence of plant OEF18 and OEF18-like clades. (A) The basal clade and outgroup taxa are highlighted in black and orange, respectively. Representative model and crop species are marked in bold text. ML bootstrap support is indicated on the individual branches; branch length is not drawn to scale. (B) Schematic comparison between the simplified ML tree topology (H1; left panel) and the null-hypothesis topology (H0; right panel) of OEF18 and OEF18L proteins across selected species.

### Calcium binding promotes homo-oligomerization of OEF18

EF-hands tend to occur in pairs, enabling co-operative binding of Ca^2+^ (Gifford *et al*., 2007). InterPro (Blum *et al*., 2025) predicted an EF-hand domain pair spanning amino acids 9-97 (IPR011992), whereas Prosite (Sigrist *et al*., 2026) identified a single EF-hand motif between amino acids 66-101 (EF_HAND_2, PS50222) in OEF18. Structural modeling of the predicted OEF18 protein structure with bound Ca²⁺ supports the presence of only one functional EF-hand fold capable of coordinating Ca²⁺ (Fig. 6A; Abramson *et al*., 2024). To determine whether OEF18 and OEF18L can bind Ca^2+^, and thus potentially alter their function in a Ca^2+^-dependent manner, we recombinantly purified full-length versions of OEF18 and OEF18L fused to glutathione-S-transferase (GST) tags. Free GST served as a negative control to exclude binding of Ca^2+^ to the purification tag, and aequorin, a Ca^2+^-sensitive bioluminescent protein with three functional EF-hands (Deng *et al*., 2005), as positive control. As expected based on EF-hand prediction, only OEF18 bound to Ca^2+^ in radio-labeled Ca^2+^ overlay assays (Fig. 6B). Furthermore, we purified the N-terminal regions of OEF18 (OEF18-N) and OEF18L (OEF18L-N) comprising the first 102 and 99 amino acids, respectively, without purification tag in a pTwin-based bacterial expression system. Additionally, a mutant version of OEF18 was generated by substituting Asp-79 in the EF-hand motif to alanine (OEF18^D79A^-N). This Ca²⁺-coordinating aspartate is evolutionarily conserved (Fig. S5) and mutation should abolish Ca²⁺-binding. Indeed, only wild-type OEF18-N showed Ca²⁺-binding ability (Fig. 6B). Bovine serum albumin (BSA) served as negative control for potential unspecific (non-EF-hand) binding of Ca²⁺ to proteins in the radio-labeled Ca^2+^ overlay assay.

**Figure 6.**
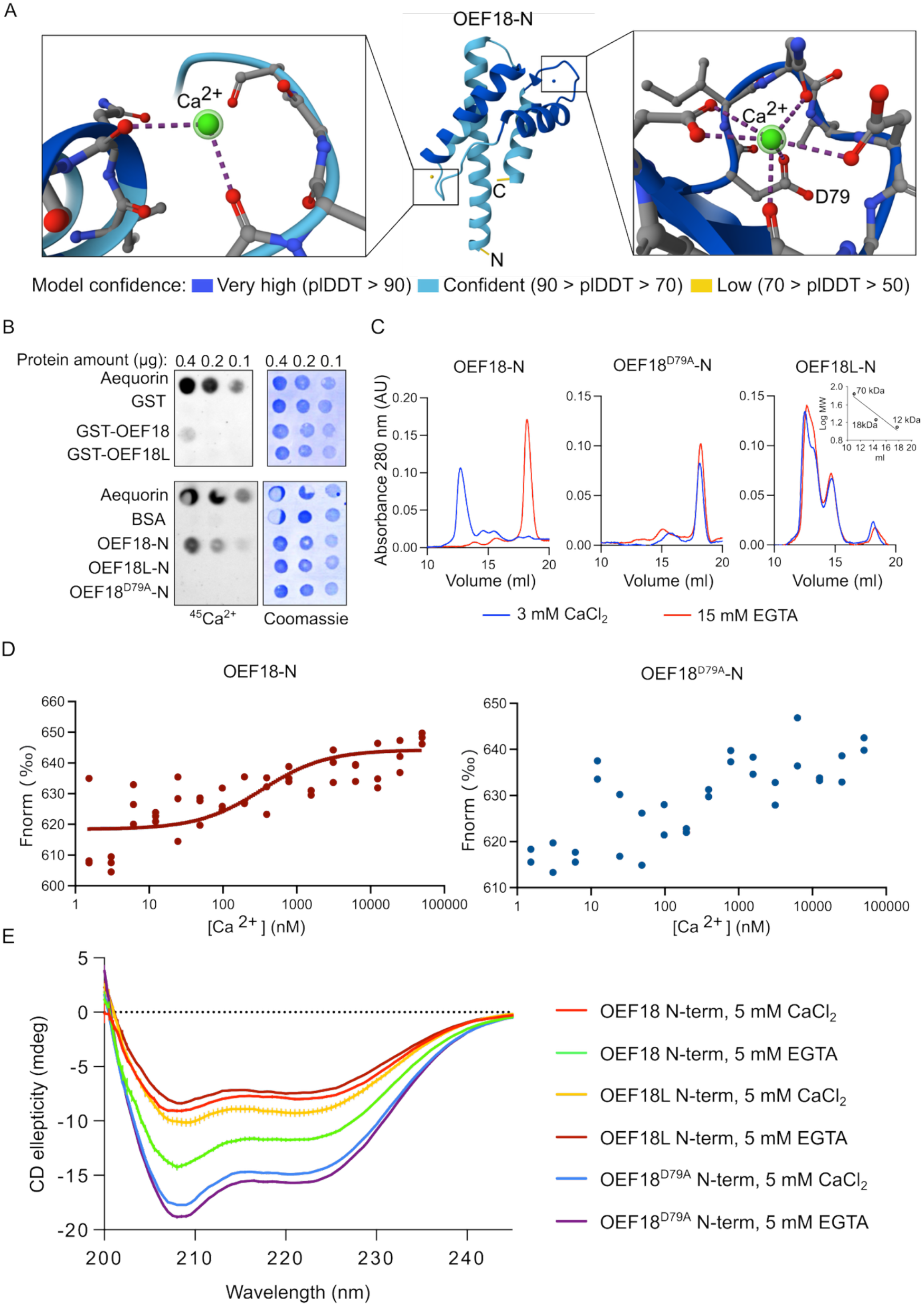
OEF18 binds Ca²⁺ and undergoes Ca²⁺-dependent oligomerization and conformational changes. (A) The structure of OEF18-N and potential Ca^2+^ binding sites predicted by AlphaFold 3. The Ca^2+^ binding site in the left panel is predicted with low confidence and lacks Ca^2+^-coordinating bonds compared to the Ca^2+^ binding site predicted with high confidence in the right panel. (B) ^45^Ca^2+^ overlay assays full-length recombinant proteins, N-terminal fragments and mutants of OEF18. Aequorin and GST or BSA served as positive and negative controls, respectively. (C) Size exclusion chromatography of OEF18-N, OEF18^D79A^-N and OEF18L-N in the presence of Ca²⁺ or EGTA. (D) MST analysis of Ca²⁺-binding affinity of OEF18-N (left panel) and OEF18^D79A^-N (right panel). F_Norm_ [‰] indicates normalized fluorescence per mill. (E) CD spectroscopy of OEF18-N, OEF18^D79A^-N and OEF18L-N in the presence of Ca²⁺ or EGTA.

To test if OEF18 might form EF-hand pairs through homo-dimerization, we subjected the N-terminal purified proteins to size exclusion chromatography in the presence of 3 mM CaCl_2_ or in the presence of a five-time excess of the Ca²⁺ chelator EGTA (ethylene glycol-bis (β-aminoethyl ether)-N,N,N′,N′-tetra acetic acid). Based on elution times compared to a calibration curve of proteins with known molecular weights, OEF18-N formed tetramers in the presence of Ca²⁺ and monomers in its absence after EGTA-mediated Ca²⁺ chelation (Fig. 6C). This Ca²⁺-dependent oligomerization was lost in the OEF18^D79A^-N mutant. In contrast, OEF18L-N formed a mixture of tetramers, dimers, and monomers irrespective of Ca²⁺ availability. Apparent molecular weight depends on the conformation of the oligomer, so without structural information on the oligomer only rough estimates are possible. Further supporting oligomerization though, OEF18 and OEF18L were found to make homo- and heterodimers at the chloroplast envelope through a transient BiFC assay in tobacco (Suppl. Fig. 7C-D). By circular dichroism (CD) spectroscopy in the far-UV range (200-240 nm), OEF18-N protein exhibited a Ca²⁺-dependent alteration of its spectral profile, indicative of a conformational rearrangement and altered secondary structure. In contrast, OEF18L-N showed only minor changes in the CD spectrum upon Ca²⁺ addition, suggesting a substantially weaker structural response to Ca²⁺ binding (Fig. 6E). The spectra revealed Ca²⁺-dependent changes in ellipticity in OEF18-N, indicating an increase in α-helical content. The OEF18^D79A^-N mutant exhibited negligible spectral differences with or without Ca²⁺, reinforcing the importance of Asp-79 for Ca²⁺-binding and resulting structural rearrangements of the OEF18 protein. Moreover, we measured the Ca²⁺-binding affinity of OEF18-N by micro scale thermophoresis. OEF18-N displayed a dose-dependent increase in normalized fluorescence with a sigmoidal profile, consistent with Ca²⁺ binding, and an apparent dissociation constant (Kd) in the range of 339.9 +/- 6.06 nM (Fig. 6D). Hill curve fitting failed for OEF18^D79A^-N, indicating a lack of strong Ca²⁺ affinity. Together, these data support the notion that OEF18 can respond to physiological levels of Ca²⁺ in the cytosol, potentially through tetramer formation, consistent with a role for OEF18 in calcium signaling.

### Accumulation of JA and related metabolites is altered in oef18 mutants, leading to reduced resistance to caterpillar herbivory

As OEF18 protein is strongly accumulated in senescent leaves (Fig. 4D), we subjected *oef18*, *oef18l* and o*ef18 oef18l* double mutants to dark-induced senescence. However, no significant difference could be observed between mutant and wild type plants after five or ten days of dark-induced senescence treatment (Fig. S8). To investigate alternative physiological roles of OEF18, we considered a potential link to JA hormone signaling. Because JA accumulation promotes senescence (He *et al*., 2002) and biosynthesis of its precursor involves metabolic steps in both chloroplasts and peroxisomes (Wasternack and Song, 2017), we examined whether loss of OEF18 affects JA-responsive gene expression. Key JA marker genes, including *LIPOXYGENASE 2 (LOX2), CHLOROPHYLLASE 1 (CHL1), PLANT DEFENSIN 1.2 (PDF1.2), and VEGETATIVE STORAGE PROTEIN 1 (VSP1)* were tested in the single *oef18* and *oef18l* mutants by RT-PCR. *LOX2*, *CHL1*, and *PDF1.2* expression was reduced in two *oef18* mutant alleles compared to wild type and *oef18l* mutants grown on soil for 4 weeks in steady-state conditions (Fig. 7A). In contrast, marker gene expression for other major plant hormone pathways, including ethylene (ET), auxin (IAA), abscisic acid (ABA), and brassinosteroids (BR), were unaffected. Interestingly, expression of the salicylic acid (SA) marker gene *PR1 (PATHOGENESIS RELATED 1)* was also down-regulated in *oef18-1* and *oef18-2*.

**Figure 7.**
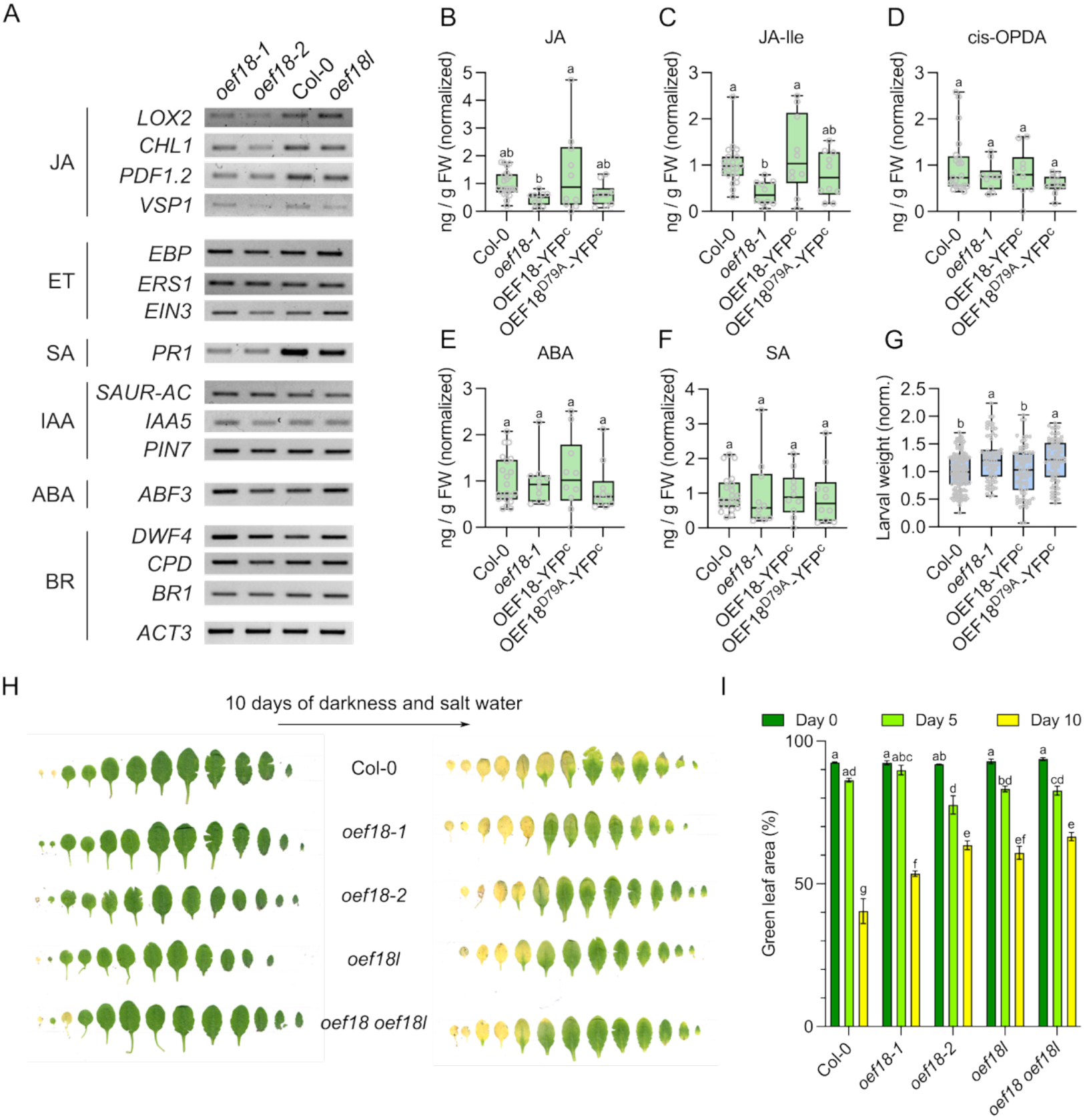
OEF18 modulates JA signaling, herbivore resistance, and combined darkness-salinity stress. (A) Semi-quantitative RT-PCR analysis of marker genes associated with different hormone biosynthesis and signaling pathways in *oef18-1*, *oef18-2*, wild type (Col-0), and *oef18l* plants. ACT3 was used as an internal control to normalize the template amount used for the reactions. (B-F) Concentration of phytohormones and related metabolites in OEF18 lines fed for 3 hours to *Spodoptera littoralis* larvae. Different letters indicate statistically significant differences (P < 0.05, one-way ANOVA with Tukey’s test). (G) Larval weight of *Spodoptera littoralis* after feeding 7 days (normalized to initial weight) on the indicated plant genotypes. Different letters indicate statistically significant differences (P < 0.05, one-way ANOVA with Tukey’s test). (H) Rosette leaf series from indicated genotypes at different developmental stages after combined darkness and salt treatment. (I) Quantification of green leaf area (%) of the indicated genotypes after combined darkness and salt treatment. Data represent mean ± SD. Different letters indicate statistically significant differences (P < 0.05, two-way ANOVA with Tukey’s test).

The contrast in altered JA marker gene expression and the lack of a dark-induced senescence phenotype prompted us to investigate another major JA-dependent stress response: caterpillar herbivory and resistance. Consistent with a role for OEF18 in JA metabolism, JA accumulation was reduced in the *oef18-1* mutant, while it was restored to wild type levels in the OEF18-YFP^c^ line after three hours of feeding by fourth instar *Spodoptera littoralis* caterpillars (Fig. 7B). Isoleucine-conjugated JA (JA-Ile), the bioactive jasmonate responsible for activating JA signaling through the COI1 receptor complex (Fonseca *et al*., 2009; Sheard *et al*., 2010), exhibited changes that mirrored those observed for JA itself (Fig. 7C). Similar trends were observed for downstream JA catabolites, including COOH-JA-Ile, OH-JA-Ile, and OH-JA (Fig. S9A-C), indicating that alterations in JA homeostasis extend throughout the JA metabolic pathway. In contrast, accumulation of 12-oxo-phytodienoic acid (OPDA), the chloroplast-derived precursor of JA that is converted to JA following transport to the peroxisome and β-oxidation, was unaffected (Fig. 7D). These results suggest that OEF18 influences JA metabolism downstream of OPDA biosynthesis, affecting JA accumulation and turnover rather than the production of its immediate precursor. ABA and SA hormone levels were similarly unaffected (Fig. 7E and 7F). In contrast to the marker gene expression data, JA and related metabolites were unaltered during steady-state conditions when grown on soil (Fig. S9D-K).

Finally, to assess the biological consequences of altered JA levels, we evaluated the performance of *S. littoralis* caterpillars on *oef18* mutant plants. First instar larvae feeding on *oef18-1* mutants gained more weight than those on wild type OEF18-YFP^c^ plants (Fig. 7G), consistent with a reduction in the levels of JA and its metabolites in the *oef18-1* plants. Importantly, a complementation line in the *oef18-1* background in which the EF-hand of OEF18 was mutated (OEF18^D79A^-YFP^c^) was unable to rescue this larval weight phenotype (Fig. 7G), which was reflected in the reduced accumulation of JA and its metabolites in the OEF18^D79A^-YFP^c^ plants (Fig. 7B-D and Fig. S9D-I). These results indicate that OEF18 positively regulates JA accumulation during caterpillar herbivory. Loss of OEF18 reduced this response and enhanced herbivore performance, and this function depended on an intact Ca²⁺-binding EF-hand motif, as the OEF18^D79A^-YFP^c^ line failed to restore JA levels or caterpillar resistance.

## Discussion

Our initial aim to identify putative EF-hand containing proteins within the chloroplast stroma led to the discovery of OEF18. Contrary to our expectation of identifying stromal or thylakoid-associated EF-hand proteins, OEF18 was instead localized to the chloroplast outer envelope, with its functional EF-hand domain exposed to the cytosol. Previously, we identified another EF-hand protein associated with the chloroplast envelope, a mitochondrial carrier family protein named S-adenosylmethionine transporter-like (SAMTL). SAMTL is likely localized to the inner envelope membrane and has been proposed to mediate the import of S-adenosylmethionine (SAM) into the chloroplast in a Ca²⁺-dependent manner (Stael, Rocha, *et al*., 2011). Consequently, the stroma and thylakoids appear largely devoid of canonical EF-hand proteins, with CRSH as the only well-characterized example reported in rice and *A. thaliana* (Masuda *et al.,* 2008; Tozawa *et al*., 2007). This may reflect the atypical Ca²⁺ dynamics of the stroma, characterized by prolonged Ca²⁺ transients during light-dark transitions (Sai and Johnson, 2002) as well as in response to high light and heat stress (Lenzoni and Knight, 2019; Kuang *et al*., 2025). Despite this unexpected localization, our findings reveal that OEF18 function influences JA accumulation, a key phytohormone whose biosynthesis is initiated in the chloroplast stroma and is central to herbivore defense. Importantly, this function appears to be Ca²⁺-dependent, as mutation of the EF-hand in the *A. thaliana* OEF18^D79A^-YFP^c^ line reduced both JA accumulation and complementation of the herbivore resistance phenotype.

A detailed study of the localization revealed curious inter-chloroplast OEF18-YFP accumulations (Fig. 1F-I). A similar localization pattern has been reported for At4g27610, a tail-anchored protein of unknown function that localizes to the chloroplast envelope (Lee *et al*., 2011). However, the apparent enrichment at chloroplast-chloroplast interfaces was not discussed by the authors and overall inter-chloroplast protein localization seems an understudied phenomenon. Chloroplasts are not symmetrically round, but can be described as flattened ovoid concave disks (Harwood *et al*., 2020). OEF18-YFP was found at the rim or belt of this disk in free floating mesophyll cells (Fig. 1D), and it is tempting to speculate that upon prolonged contact between chloroplasts, OEF18-YFP might accumulate at the interface of clustered chloroplasts (Fig. 1F-I), thereby enlarging the surface interaction of adjacent chloroplasts. As OEF18 can homo-oligomerize in a Ca²⁺-dependent manner (Fig. 6C) it is possible that OEF18 is required for this tight chloroplast-to-chloroplast interaction. Alternatively, OEF18 is not required for interaction and rather modulates a particular function at this site. A potential caveat regarding our observations of chloroplast appressions is that fluorescent protein fusions can introduce artificial structural phenotypes, as the YFP variant used in this study (eYFP) is known to form a weak dimer. However, potential YFP dimerization does not necessarily explain the preferred accumulation of the OEF18-YFP fusion protein at the chloroplast interface. Several lines of evidence from native, label-free ultrastructural studies indicate that the tight chloroplast contacts observed here are unlikely to be transgenic artifacts. Volumetric reconstructions of wild-type mesophyll tissue by Serial Block-Face SEM (SBF-SEM) and Focused Ion Beam SEM (FIB-SEM) show that chloroplasts are naturally irregularly shaped and frequently crowded against one another (Knoblauch *et al*., 2024; Harwood *et al*., 2020; Yamane *et al*., 2026). The function of these closely appressed chloroplast interaction surfaces remains unclear. One possibility is that they facilitate the exchange of metabolites or signaling molecules, including Ca²⁺ or intermediates of the JA pathway. Alternatively, such associations may promote more efficient spatial organization of chloroplasts within the cell, minimizing unoccupied cytoplasmic space and maximizing the photosynthetic surface area exposed to incoming light, thereby enhancing light capture efficiency. However, these potential functions of OEF18 and chloroplast appressions remain highly speculative and will require further investigation.

OEF18 displays unique features that influence its subcellular localization, while also sharing some characteristics with TA proteins. In plants, TA proteins are defined by a single C-terminal transmembrane domain, a cytosolic N-terminus, and the absence of an N-terminal signal peptide, being inserted post-translationally into target membranes such as the ER, mitochondria, and chloroplasts. Unlike typical TA proteins, OEF18 contains two transmembrane domains, but similar to TA proteins, OEF18 interacts with the cytosolic chaperone AKR2A, which probably aids in membrane targeting of OEF18 by recognizing hydrophobic domains and charged residues near the C-terminus (Fig. 3A-C). TA proteins can exhibit dual or multiple subcellular localizations. For instance, FISSION1A has been shown to target chloroplasts, peroxisomes, and mitochondria (Ruberti *et al*., 2014). Evidence for peroxisomal localization of OEF18 is mixed: it is absent in the stable native promoter OEF18-YFP lines (Fig. S1B), but consistently observed in transient expression assays in tobacco and by sucrose density gradient fractionation in *A. thaliana* (Fig. 1B and 1J). Thus, a peroxisomal localization cannot be totally excluded. In contrast, chloroplast envelope localization was consistently detected across all experimental approaches. Taken together, myristoylation and the transmembrane domain are crucial determinants for targeting of OEF18 to the chloroplast outer envelope and potentially to the peroxisome membrane.

The localization of OEF18 at the chloroplast outer envelope, with its EF-hand domain exposed to the cytosol, places it to potentially sense or relay Ca²⁺ signals originating outside the organelle. Recent studies have highlighted the importance of ER-chloroplast membrane contact sites as hubs for inter-organelle communication, including the exchanges of biomolecules such as lipids, proteins, hormones and Ca²⁺ signals (Depaepe and Munné-Bosch, 2026). As one of the major intracellular Ca²⁺ stores in plant cells, the ER acts as a vital buffering and releasing system to maintain cellular Ca²⁺ homeostasis (Grenzi *et al*., 2026). Consistent with this, our previous work has shown that high light-induced stromal Ca²⁺ transients are mirrored by the ER Ca^2+^ pool, indicating potential ER-chloroplast interactions in plant cells during the high light response (Kuang *et al*., 2025). Previously, a chloroplast-localized mitochondrial calcium uniporter (cMCU) was described that transduces osmotic stress signals into chloroplasts (Teardo *et al*., 2019). In addition, with the identification of Ca²⁺ transporters such as BIVALENT CATION TRANSPORTER 2 (BICAT2) (Frank *et al*., 2019) and the involvement of PLASTID ENVELOPE ION CHANNELS (PEC1/2) on stress-induced stromal Ca^2+^ dynamics (Völkner *et al*., 2021), the chloroplast envelope has emerged as an important interface for the integration of Ca²⁺ dynamics. Together, these findings suggest that coordinated Ca²⁺ exchange between these organelles may contribute to stress signaling, raising the possibility that OEF18 may couple cytosolic or ER-derived Ca²⁺ dynamics to chloroplast-associated metabolic and defense responses. Although this hypothesis awaits direct experimental validation, it is consistent with the emerging concept that ER-chloroplast membrane contact sites enable rapid and spatially restricted Ca²⁺ signaling between organelles.

During the course of our study, the maize ortholog of OEF18, named ZmNSA1 (*Zea mays* Na^+^ Content under Saline-Alkaline Condition; GRMZM2G000397), was found to confer tolerance to saline-alkaline stress (Cao *et al*., 2020). ZmNSA1 was identified through a genome-wide association study of maize inbred lines grown on NaHCO_3_, revealing a 4-bp deletion in its promoter that reduces ZmNSA1 protein amount and decreased Na⁺ accumulation in the shoot. ZmNSA1 was predicted to contain EF-hand motifs, and its Ca²⁺-binding affinity (Kd ≈ 220 nM) is in the same sub-micromolar range as that determined for OEF18 (Kd ≈ 340 nM). Based on transient expression in maize protoplasts of a ZmNSA1-GFP fusion protein, the authors suggest that ZmNSA1 is primarily localized in the cytosol. However, the presence of distinct rings around chloroplasts, which may indicate additional localization, was not discussed, nor were other localization studies performed. Considering the strong sequence conservation of OEF18 orthologs and their transmembrane domains (Fig. S5), we suggest that ZmNSA1 is likely not localized to the cytosol but rather to the chloroplast envelope, and potentially the peroxisome membrane. Consistent with previous findings, we observed OEF18 protein accumulation is slightly enhanced in osmotically and salt-stressed seedlings treated with 100 mM Sorbitol or 75 mM NaCl, respectively (Fig. S8A). OEF18 protein levels were markedly increased during natural senescence (Fig. 4D). Consequently, a combination of dark-induced senescence and salt stress led to delayed senescence symptoms in *oef18* and *oef18l* mutants (Fig. 7H and I). This may reflect higher Na⁺ accumulation and toxicity in WT plants. Interestingly, ZmNSA1 protein levels were found to be regulated post-transcriptionally, as the proteasome inhibitor MG132 suppressed protein degradation during NaHCO₃ treatment, with higher ZmNSA1 levels correlating to higher Na⁺ accumulation. Together, these findings support the conclusion that reduced OEF18 levels correlate with enhanced Na⁺ tolerance.

Cao *et al*. (2020) further conclude that under saline-alkaline conditions, Ca²⁺-triggered degradation of ZmNSA1 promotes the transcriptional upregulation of plasma membrane H⁺-ATPases MHA2 and MHA4, leading to enhanced root H⁺ efflux. This activation of H⁺-ATPase activity supports SOS1-mediated Na⁺ efflux and contributes to improved salt tolerance in maize. However, it remained unclear how the amount of ZmNSA1 protein is connected to the transcriptional regulation of plasma membrane H⁺-ATPase. Previous work demonstrates a link between JA signaling and plasma membrane proton pump activity, particularly at the transcriptional level. Studies in *Cucumis sativus* provided evidence of JA-mediated transcriptional downregulation of plasma membrane H⁺-ATPases *CsHA1, CsHA3*, and *CsHA9* (Janicka *et al*., 2023). However, the relationship is likely more complex as a feedback loop exists, exemplified by *A. thaliana* AHA1, which modulates JA signaling, primarily through its influence on membrane potential and electrical signals (Kumari *et al*., 2019). The molecular mechanisms by which core JA components, such as MYC2, regulate plasma membrane H⁺-ATPase gene transcription remain unclear, and the potential role of OEF18 in this regulatory pathway also awaits further elucidation. Finally, while reducing OEF18 accumulation using CRISPR/Cas technology could enhance saline-alkaline stress tolerance in crop species, potential trade-offs, such as altered JA metabolism and increased susceptibility to herbivores, should be carefully evaluated.

Altogether, we describe here a novel type of EF-hand protein at the outer chloroplast envelope that connects chloroplasts and potentially peroxisomes with their environment, i.e. the cytosol or other organelles and suggest that it mediates organellar interactions in response to external stimuli generating Ca^2+^ signals.

## Limitations

Despite the accumulated data from *A. thaliana* and *Zea mays*, the mechanistic role of OEF18 remains unresolved. In particular, it is unclear how a Ca²⁺-binding protein at the chloroplast outer envelope regulates JA metabolism, and whether this reflects a direct regulatory function in JA biosynthesis or an indirect consequence of altered chloroplast Ca²⁺ signaling or organelle organization. Interestingly in this respect is that chloroplast morphology seems to affect JA metabolism (Baral *et al*., 2026). Future work could therefore combine inducible perturbation of chloroplast morphology with quantitative JA measurements to determine whether altered plastid architecture is sufficient to account for the metabolic phenotypes observed in *oef18* mutants. Likewise, the proposed involvement of OEF18 in chloroplast-chloroplast contact sites and inter-organelle interactions remains speculative, as does the significance of its occasional peroxisomal association. Future work could address this by inducible induction of chloroplast clustering for example with knock sideways technology (Winkler *et al*., 2021). Finally, it is not yet clear whether Ca²⁺-dependent oligomerization of OEF18 represents a functional signaling mechanism in vivo or a structural property linked to membrane clustering, leaving the primary mode of action of OEF18 unresolved. Future work should therefore combine separation-of-function mutants (Ca²⁺ binding and oligomerization), stable organelle-specific complementation, and proximity labelling approaches to define OEF18 interaction networks at the chloroplast envelope. In parallel, compartment-resolved Ca²⁺ imaging and targeted JA pathway readouts will be required to directly link envelope Ca²⁺ dynamics to metabolic outputs and clarify whether OEF18 acts as a true Ca²⁺-dependent signaling hub or reflects broader changes in chloroplast organization.

## Material and methods

### Plant material and growth conditions

For media-grown plants, *A. thaliana* seeds were surface-sterilized with 70% (v/v) ethanol for 20 min followed by 95% (v/v) ethanol for 5 min, and subsequently stratified in sterile distilled water at 4 °C for 2 d. Seedlings were grown vertically on half-strength Murashige and Skoog (½MS) medium, supplemented with 0.8% (w/v) agar and 0.5% (w/v) sucrose for 12 days at 22 °C/ 20 °C with a day/night cycle of 16/8 h.

For soil-grown plants, *A. thaliana* seeds were sown on a soil mixture composed of four parts Huminsubstrat N3 (Neuhaus) and one part Perlite (Gramoflor), supplemented with Osmocote (Substral/Scotts) according to the manufacturer’s instructions. Seeds were stratified at 4 ℃ for 2 d and then transferred to the appropriate growth chambers under the conditions specified for each experiment. For stress assays, seedlings were grown on 1/2 MS medium containing either 1 μL/L paraquat, 100 mM sorbitol, or 75 mM NaCl. Seedlings maintained on standard 1/2 MS medium were used as controls. After 2 weeks of treatment, whole seedlings were harvested, immediately frozen in liquid nitrogen and stored at -80 °C until further analysis. Total protein was extracted from frozen tissues using 2× laemmli buffer and analyzed by immunoblotting as described in the Western blot section.

### Transient expression in Nicotiana tabacum leaves

For co-localization, BiFC, and saGFP assays, *Agrobacterium tumefaciens* strain AGL1 carrying the respective constructs was cultured overnight at 30 °C in 5 ml LB medium containing kanamycin. This culture was subsequently expanded in 50 ml LB medium with kanamycin and incubated for 3 h at 30°C. The bacteria were precipitated by centrifugation at 3500 rpm for 15 min at room temperature and resuspended in 30ml LB medium supplemented with 100mM acetosyringone, followed by incubated for 2 h at 30°C. The bacteria were centrifuged again and resuspended in a 5% (w/v) sucrose solution to an optical density (OD600) of approximately 2.0. Leaf infiltration was performed using a needleless syringe by gently pressing the suspension onto the underside of leaves of 4-week-old *Nicotiana tabacum* plants. Successful infiltration was indicated by the appearance of a spreading “wetting” area in the leaf. For co-localization, BiFC or saGFP assays, equal volumes of the respective bacterial suspensions were mixed before infiltration. After infiltration, plants were covered with plastic bags to keep high humidity. Covers were removed after 24 h and samples were used for microscopy analysis the following day.

### Generation of OEF18 complementation lines

A genomic fragment encompassing approximately 1 kb upstream of the OEF18 translational start codon (and extending to the native stop codon, including all exons and introns, was amplified by PCR from *A. thaliana* genomic DNA and cloned using ApaI and NotI restriction sites into the vector pBIN-Basta, a derivate of pBIN 19 (Bevan, 1984) carrying a C-terminal YFP fusion. The resulting constructs were introduced into *A. tumefaciens* strain GV3101 for plant transformation, and transgenic plants were selected on medium containing Basta (glufosinate ammonium). T2 transformants were subjected to Mendelian segregation analysis to identify single-copy insertion lines, and homozygous T3 lines were selected for further analysis. Primer sequences used for cloning are listed in Table S2. The resulting pOEF18::OEF18-YFP/*oef18-1* line, abbreviated as OEF18-YFP^c,^ was further transformed with a peroxisomal marker, mScarlet-SKL driven by an *A. thaliana* UBIQUITIN10 promoter and terminator (AtUBI10p:mScarlet-SKL:AtUBI10t).

### Confocal microscopy

*N. tabacum* leaf samples were infiltrated with distilled water using a 50 mL syringe (Braun) to improve optical properties of the sample. Confocal laser scanning microscopy (CLSM) was performed on a Leica confocal microscope. The samples were mounted on glass slides and secured with cover slips using double-sided tape. Images were acquired with a 20× 0.75 NA dry objective.

Brightfield and fluorescence channels were acquired simultaneously with the following settings: CFP (Ex: 405 nm, 30% power; Em: 460-490 nm), GFP (Ex: 488 nm, 30% power; Em: 500-525 nm), YFP (Ex: 514 nm, 30% power; Em: 520-550 nm), mCherry (Ex: 587 nm, 70% power; Em: 600-615 nm), and chlorophyll autofluorescence (Ex: 514 nm, 30% power; Em: 630-750 nm). Obtained images were analyzed using ImageJ software.

CLSM of *A. thaliana* seedlings was performed on a Nikon AX R confocal microscope. Cotyledon samples from 10-12-day-old *A. thaliana* seedlings (OEF18-YFP^c^ and OEF18-YFP^c^ × mScarlet-SKL double line) were detached and placed on glass slides with distilled water and secured with cover slips. Images were acquired at 512 × 512 pixels, with a 40× 1.25 NA silicone immersion objective and 3X digital zoom. Brightfield and fluorescence channels were acquired simultaneously with the following settings: YFP (Ex: 514 nm, 3% power; Em: 520-535 nm), mScarlet (Ex: 561 nm, 0.5% power; Em: 580-630 nm), and chlorophyll autofluorescence (Ex: 640 nm, 1% power; Em: 662-737 nm). Advanced noise removal was performed using the trained AI deep-learning algorithm (Nikon NIS-Elements Denoise.AI). Images were further analyzed using ImageJ software (Schindelin *et al*., 2012) .

### Fluorescence intensity quantification of OEF18 inter-chloroplast accmulation

Quantitative fluorescence intensity was analyzed on the YFP channel using ImageJ software. For each target chloroplast, regions of interest (ROIs) were manually selected using the polygon selection tool at inter-chloroplast accumulations and the adjacent envelope periphery. Mean gray values were extracted and normalized to the average envelope intensity using GraphPad Prism.

### Chloroplast isolation and sub fractionation

Chloroplasts were isolated from 4-week-old *A. thaliana* plants grown under short-day conditions, according to (Bayer *et al*., 2011) with modifications. Leaf tissue (80-100g) was homogenized in grinding buffer (0.4M sorbitol, 20mM Tricine, 10mM Na_2_EDTA, 5mM NaHCO_3_, 0.1% BSA, 10 mM ascorbic acid, pH 8.4 adjusted with KOH) at a ratio of 1:4 (w/v). All following steps were carried out on ice or at 4 °C. The homogenate was filtered through triple layers of Miracloth (Merck Millipore) and centrifuged at 2700 rpm for 2 min. The organelle pellet was then gently resuspended in 5ml washing buffer (0.4M sorbitol, 20mM Tricine, 10mM MgCl_2_, 5mM Na_2_EDTA, pH 7.6 adjusted with KOH) and loaded onto the 40%-80% Percoll gradients. These gradients were then centrifuged at 4000 rpm for 20 min in swing-out rotor with low deceleration. Intact chloroplasts at the interface of the gradient were collected and washed twice with 50ml washing buffer. The final chloroplast pellet was resuspended in 500 µl washing buffer. The quantity of chloroplast was assessed by the amount of chlorophyll determined based on Porra’s method (Porra *et al*., 1989). The quality of purified chloroplast was analysed by light microscopy. The isolated chloroplasts were used immediately or stored at -80 °C for further experiments.

To fractionate chloroplasts into thylakoid, stroma and envelope, Percoll-purified chloroplasts corresponding to 1 mg of chlorophyll were resuspended in cold Tris-EDTA (TE) buffer and lysed on ice for 10 min. Lysed chloroplasts were overlaid on a discontinuous sucrose density gradient (1.2 M, 1.0 M, 0.46 M in TE buffer) and centrifuged at 18000 rpm for 2 h at 4 °C in a swinging-bucket rotor. The top fraction corresponding to stroma was precipitated by chloroform-methanol method. The intermediate envelope fraction and the thylakoid fraction at the bottom were collected separately, both resuspended in 2 mL of TE buffer, and centrifuged at 30000 rpm for 1 h and 4000 rpm for 5 min at 4 °C, respectively. Fractions were resuspended in TE buffer followed by western blot.

### Fractionation of A. thaliana membranes by sucrose density gradient centrifugation

The experiment was performed according to (Lu and Hrabak, 2002). All steps were performed at 4°C. Approximately 10-12 g of *A. thaliana* leaves were collected from 8-week-old plants for each gradient. The leaves were thoroughly homogenized in 100 ml homogenization buffer (50 mM Tris-HCl (pH 7.8), 400 mM sorbitol, 2 mM EDTA, 15 mM MgCl_2_, 1 mM DTT and cOmplete, EDTA-free Protease Inhibitor Cocktail) using a Waring blender (5 pulses of 3 s at maximum speed). The homogenate was filtered through double layers of Miracloth (Merck) and centrifuged at 10.000 × g for 20 min in a SW28 rotor (Beckman Coulter) at 4 °C to remove most large debris, including nuclei and thylakoids. The supernatant was subsequently centrifuged at 100.000 × g for 1 h in the same rotor to precipitate the microsomal fraction. The microsomal pellet was rinsed with microsome resuspension buffer (10 mM Tris-HCl (pH 7.5), 2 mM EDTA, 15 mM MgCl_2_, 55% (w/w) sucrose) and mechanically disrupted. The suspension volume was adjusted to 0.5 ml and carefully transferred to the bottom of a 14 ml thin-wall polyallomer centrifuge tube. A linear sucrose gradient (20-50% w/w) was generated using a gradient mixer and layered over the sample (total volume 6 ml). The gradients were centrifuged at 100.000 × g for 18 h at 4 °C in an SW40Ti rotor (Beckman Coulter). Fractions were collected sequentially from the top of the gradient and the sucrose concentrations in each fraction was measured with a refractometer (Pal-1, Atago). Fractions were stored at -20°C until further analysis.

### Calcium overlay assay

Radioactive ^45^Ca^2+^ overlay assays were performed according to Maruyama *et al*., (1984), with modifications. Dilution series of purified protein (5, 2.5 and 1.25 µg) were spotted on activated PVDF membrane (Merck Millipore), after which membranes were then placed in fixing solution containing 10% (v/v) isopropanol and 10% (v/v) acetic acid, followed by washing in the assay buffer (60mM KCl, 5mM MgCl2, 10mM imidazole pH 6.8). Membranes were incubated in the assay buffer supplemented with 1µM ^45^CaCl_2_ (1µCi/ml; PerkinElmer) for 15 min and washed 4 times for 30 s with deionized water, air-dried and exposed to a Storage Phosphor Screen (GE Healthcare) overnight. Signals were detected using a Typhoon 8600 Variable Mode Imager (Amersham). Protein loading was subsequently verified by Coomassie Brilliant Blue staining.

### Gel filtration chromatography

Size-exclusion chromatography was performed using a Superdex 75 10/300 column (GE Healthcare) connected to an FPLC system (GE Healthcare). The column was washed with at least 3 column volumes (CV; 1 CV=24 ml) of distilled water and then equilibrated with 2 CV of assay buffer (20 mM HEPES, pH 7.5, 10 mM MgCl₂) supplemented with 3 mM CaCl₂ or 15 mM EGTA. Column calibration was performed using 300µl standard protein mixture (cytochrome c, lysozyme and BSA). Samples were loaded into a 500 µl sample loop. A calibration curve was generated by plotting the log of molecular weight against the volume. For experimental samples, 500µl 0.5µg/µl protein solution in the appropriate buffer (supplemented with 3 mM CaCl₂ or 15 mM EGTA) was loaded onto the column. Flow rates were the following: 100% buffer A (assay buffer), 0% buffer B; volume 2.4ml; flow 0.8ml/min, followed by injection of the sample (dynamic loop) volume 0.6ml; flow 0.3ml/min, and protein separate afterwards with 100% buffer A (assay buffer), 0% buffer B; volume 26ml; flow 0.6ml/min. UV absorption spectra were recorded and analyzed to determine the oligomeric state of proteins.

### Western blot

Western blot was performed using a semi-dry transfer technique. SDS-PAGE-separated proteins were transferred onto PVDF membranes (Immobilon-P; Merck Millipore) at 80 mA for 50 min following equilibration in transfer buffers. Membranes were blocked in TBST buffer (50 mM Tris-HCl, pH 7.4, 150 mM NaCl, 0.1% (v/v) Tween-20) supplemented with either 1% (w/v) BSA or 5% (w/v) skimmed milk depending on the manufacturer’s suggestion for the primary antibody. Primary antibody incubation was performed overnight at 4 °C, followed by three washes in TBST. Membranes were incubated in secondary antibody for 1 h at room temperature, followed by three washes in TBST. Signals were detected using an ECL Plus Western Blotting kit (GE Healthcare) and visualized by exposure to X-ray film (Fujifilm). Films were developed using standard developer and fixer solutions (AGFA).

### Semi-quantitative RT-PCR

Frozen plant materials (leaves, whole rosettes, or roots) were ground to a fine powder in liquid nitrogen using either mortar and pestle or a Mixer Mill MM 400 (Retsch). Approximately 200 mg of tissue powder was homogenized in RNA extraction buffer (1% (w/v) SDS, 200 mM sodium acetate, 10 mM EDTA, pH 5.2) and mixed with an equal volume of acidic phenol (pH 4.0; Carl Roth). Following phase separation by centrifugation at 14.000 × g for 10 min at 4 °C for 10 min, the aqueous phase was recovered and extracted sequentially with phenol: chloroform: isoamyl alcohol (25:24:1) and chloroform. Subsequently, the aqueous phase was mixed with 1/3 volume of 10M lithium acetate and incubated overnight at 4°C, followed by centrifugation at 14.000 × g for 10 min at 4 °C. The RNA pellet was washed twice with 70% (v/v) ethanol and resuspended in RNase-free water (Sigma-Aldrich). RNA quality was assessed by electrophoresis on 1.6% agarose gel, and concentration was determined by NanoDrop (Thermo Scientific) measurement. Genomic DNA contamination was removed by treating 8 µg total RNA with RQ1 DNase (Promega) according to the manufacturer’s instructions. First-strand cDNA synthesis was performed using 2 µg total RNA, oligo(dT)₁₅ primers (Microsynth) and M-MLV reverse transcriptase (Promega) according to the manufacturer’s instructions. For RT-qPCR, M-MLV H (-) point mutant reverse transcriptase (Promega) was used.

Semi-quantitative RT-PCR was performed using DreamTaq DNA polymerase (Fermentas) according to the manufacturer’s instructions, with 1 µl cDNA as template. Actin2 was used as an internal control to normalize cDNA input across samples. The following amplification program was employed: denaturation at 95 °C for 5 min, followed by 26 cycles of 95 °C for 45 s, 55 °C for 45 s, and 72 °C for 45 s, with a final extension at 72 °C for 5 min. PCR products were separated on 1% (w/v) agarose gels and visualized by ethidium bromide staining. Annealing temperatures and cycle numbers were optimized for each primer pair to ensure amplification within the linear range. Band intensities were compared to assess relative transcript abundance between samples. Primer sequences used for RT-PCR are listed in Table S2.

### Protoplast isolation for microscopy

Protoplasts were isolated from soil-grown mature *A. thaliana* leaves by tape-sandwich method (Wu *et al*., 2009), according to Kuang *et al*., (2025).

### Insect feeding assays

*Spodoptera littoralis* (Boisduval) (Lepidoptera; Noctuidae) larvae were obtained from eggs (Syngenta Crop Protection AG, Switzerland) and reared as described previously (Müller *et al*., 2024). For long-term feeding assays, biomass measurements were performed using pre-weighed first-instar larvae to ensure equal starting conditions. Three larvae were placed on each 5-6-week-old plant, and larval weight was determined individually after 7 d of feeding. For short-term feeding assays, fourth-instar larvae were starved for 12 h prior to the experiment and allowed to feed on plants for 3 h. Following feeding, treated leaves were harvested, weighed, and flash-frozen in liquid nitrogen. Samples were stored at -80 °C for further analysis.

### Phytohormone analysis

Approximately 200 mg frozen leaf material from each plant was pooled and homogenized. Phytohormones were extracted and quantified as described by (Müller *et al*., 2022) with modifications. Briefly, samples were extracted in 1 mL of methanol containing 40 ng D_4_-SA (Santa Cruz Biotechnology, USA), 40 ng D_6_-JA (HPC Standards GmbH, Germany), 40 ng D_6_-ABA (Toronto Research Chemicals, Toronto, Canada), and 8 ng D_6_-JA-Ile (HPC Standards GmbH) as internal standards. After extraction and clarification by centrifugation, the supernatants were subjected to LC-MS/MS analysis using an Agilent 1260 series HPLC system (Agilent Technologies, Waldbronn, Germany) coupled to a tandem mass spectrometer QTRAP 6500 (SCIEX, Darmstadt, Germany). The following phytohormones were quantified: abscisic acid (ABA), 12-oxo-phytodienoic acid (cis-OPDA), jasmonic acid (JA), 12-hydroxyjasmonic acid (OH-JA), jasmonoyl-isoleucine conjugate (JA-Ile), 12-hydroxyjasmonoyl-isoleucine (OH-JA-Ile), dicarboxyjasmonoyl-isoleucine (COOH-JA-Ile), and salicylic acid (SA).

### Yeast two-hybrid assay

Positive clones selected on SD-Leu-Trp plates were cultured in 5ml liquid medium at 30 °C for 2 days. Yeast cells were harvested and lysed in lysis buffer (25 mM Tris-HCl, pH 7.5, 20 mM NaCl, 8 mM MgCl₂, 5 mM DTT, 0.1% NP-40) using glass beads and mechanical disruption. Lysates were clarified by centrifugation at 20.000 × g for 10 min at 4 °C, and the supernatant was collected for further analysis. Protein extracts were used for β-galactosidase assay to determine the strength of protein-protein interaction. Reactions were performed in 96-well plates by mixing 10 µl protein extract with 150 µl reaction mix containing Z-buffer (60 mM Na₂HPO₄, 40 mM NaH₂PO₄, 10 mM KCl, 1 mM MgSO₄, 0.5% β-mercaptoethanol) and 4 mg/ml ONPG. The reaction was incubated until a yellow color developed (up to 10 min) and terminated by adding 1 M Na₂CO₃. Absorbance was measured at 420 nm. Additionally, total protein concentration was determined using the Bradford assay. β-Galactosidase activity was calculated as following:

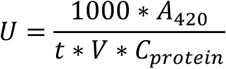

Here, ‘U’ is specific activity of β-Galactosidase (in U/mg protein); ‘A_420_’ is the absorbance of the reaction mixture at 420 nm (corrected for the blank); ‘t’ represents the incubation time (min); ‘V’ represents the volume of the cell cultures or protein extracts added for the assay (mL) mL; ‘C_protein_’ represents total protein concentration determined by the Bradford assay (mg/mL). The experiments were repeated six times.

### In vitro myristoylation assays

Analysis of protein N-myristoylation was carried out as previously described (Benetka *et al*., 2008) using a cell free system (TNT Coupled Wheat Germ Extract System, Promega). In vitro translation was carried out either in the presence of 10 μCi of L-[^35^S] methionine (1175 Ci/mmol, Perkin-Elmer) for total protein labelling, or 50 μCi of [9,10-^3^H]-labelled myristic acid (60 Ci/mmol, American Radiolabeled Chemicals). Reaction products were separated on 12% (w/v) SDS-polyacrylamide gels and incubated with autoradiography intensifier (Amersham) before detection on X-ray film.

### Genotyping and genomic DNA extraction

Genomic DNA was extracted from *A. thaliana* leaf tissue. Approximately 150 mg of frozen leaf material was homogenized in DNA extraction buffer (200mM Tris PH 7.5, 250mM NaCl, 25mM EDTA, 0.5% SDS), and cellular debris was removed by centrifugation at 20.000g for 5 min at room temperature. The supernatants were collected and precipitated with isopropanol. After another centrifugation, the pellet was washed with 70% (v/v) ethanol, air-dried, and resuspended in nuclease-free water. Genotyping was performed by PCR using gene-specific and T-DNA border primers and PCR products were analyzed by agarose gel electrophoresis. Primer sequences used for genotyping are listed in Table S2.

### Dark-induced senescence (DIS) assay and salinity treatment

DIS assays were performed as described by Mair *et al*., (2015) with modifications. *A. thaliana* plants were grown under long-day conditions for 4 weeks and subsequently transferred to a dark chamber with controlled air exchange to prevent ethylene accumulation. After 5, 8 and 10 d of dark treatment, leaves were harvested and imaged. Leaf area and green leaf area were quantified using ImageJ and the proportion of green tissue was calculated.

For combined DIS and salt stress assays, plants were treated as described above for DIS, with the addition of 200 mM NaCl applied on the day of transferring into the darkness. Data were analyzed as described for DIS.

### Circular Dichroism Spectroscopy

Circular dichroism (CD) spectroscopy was performed on recombinant OEF18 protein and its variants at 0.5 μg/μL in 10 mM HEPES buffer (pH 7.0) containing 5 mM MgCl₂. Protein samples (1 mL) were supplemented with either 5 mM CaCl₂ or EGTA and loaded into a 10 mm path-length quartz cuvette. CD spectra were recorded using a Jasco spectropolarimeter over a wavelength range of 200-245 nm. Spectra were collected from five independent measurements for each condition and averaged.

### Microscale thermophoresis Ca²⁺-binding assay

The Ca²⁺-binding affinity of OEF18-N was determined on a Monolith NT.LabelFree instrument (NanoTemper Technologies). Equal volumes of recombinant OEF18-N protein at a final concentration of 40 μM and 16 dilutions of CaCl₂ (starting from 50 mM) were mixed and incubated for 5 min at room temperature to reach binding equilibrium. Samples were then transferred into glass capillaries and measured at an MST power of 40 % and an excitation power of 20 %. Fluorescence was measured before laser heating (F_Initial_) and after 20 s of laser on time (F_Hot_). The normalized fluorescence F_Norm_= F_Hot_/F_Initial_ reflects the concentration ratio of labeled molecules. F_norm_ was plotted directly and multiplied by a factor of 10, yielding a relative change in fluorescence per mill. Kd was calculated from three independent thermophoresis measurements using NanoTemper Software (NanoTemper Technologies).

### Venn diagram analysis

Protein localization datasets for chloroplasts and peroxisomes were obtained from the Plant Proteomics Database (PPDB; (Sun *et al*., 2009)), and subcellular localization was further verified using the SUBA5 database (Hooper *et al*., 2017). The list of tail-anchored (TA) proteins was obtained from Brito *et al*., (2019), and experimentally identified myristoylated proteins were retrieved from Majeran *et al*., (2018). Overlaps between TA proteins, myristoylated proteins, and organelle-localized protein datasets were identified and visualized using Venn diagrams provided by University of Ghent (http://bioinformatics.psb.ugent.be/webtools/Venn/).

### Targeted OEF18 and OEF18L orthologs search

We used the OEF18 sequences of *A. thaliana* (UniProtID: Q9XIR0) and of *Micromonas pusilla* (NCBI Accession: XP_002500805), and the OEF18L sequence from *A. thaliana* (UniProtID: A0A1P8B468) as seeds for a targeted ortholog search with fDOG (Tran *et al*., 2025) in a collection of 1101 eukaryotic genome assemblies (Muelbaier *et al*., 2024) supplemented with gene sets from three streptophyte algae obtained from GenomeZoo / MAdLandDB (https://github.com/Rensing-Lab/Genome-Zoo): *Chara braunii*, *Chaetosphaeridium globosum*, and *Chlorokybus atmophyticus*. fDOG was run with default parameters setting the parameter *minDist* to ‘class’ and *maxDist* to ‘kingdom’ during core ortholog compilation. The resulting phylogenetic profiles were visualized and analyzed with PhyloProfile v2.6.0 (Tran *et al*., 2018; https://www.github.com/BIONF/phyloprofile).

### Phylogenetic analysis and multiple sequence alignment

Orthologous sequences for OEF18 and OEF18L from the Viridiplantae were extracted from the phylogenetic profiles using for each species the representative orthologs as identified by dDOG. The sequence length of OEF18 across the viridplantae is conserved with the *A. thaliana* and *M. pusilla* orthologs differing only by 3 amino acids (162 aa vs. 165 aa). To reduce the impact of erroneous gene models on the phylogenetic analysis, we filtered out orthologs with a length of more than 1.5 x that of the *A. thaliana* sequence, setting the inclusion cut-off to 250 aa. Orthologous sequences were aligned with Muscle v5.3 (Edgar, 2004) and alignment positions with more than 50% gaps were removed with a custom perl script. An ML tree together with 1000 ultra-fast bootstrap replicates was computed from the trimmed alignment with IQ-TREE (Minh *et al*., 2020) letting the software automatically determine the best-fitting substitution model (Kalyaanamoorthy *et al*., 2017). The ML tree was visualized and processed with iTOL (Letunic and Bork, 2024). Alternative tree topologies were created with phylomatt (https://www.github.com/bionf/phylomatt) and tested against the ML tree using the AU test (Shimodaira, 2002) implemented into phylomatt.

For the multiple sequence alignment of OEF18, orthologs were retrieved from OrthoDB (Tegenfeldt *et al*., 2025) and sequences that exceeded the median protein length of 171 aa +/-20 amino acids were removed. Sequences were aligned using Clustal Omega (Sievers and Higgins, 2018) and visualized with JalView (Waterhouse *et al*., 2009).

## Acknowledgements

This work has been funded by the Austria Science Fund (FWF) (projects P28491-B29 and P37245-B) to BW and MT; the Knut and Alice Wallenberg Foundation (grant 2021.0071), the European Research Council (Consolidator grant 101044878) and Vetenskapsrådet (grant 2025-05378) to SS. Additional support was provided by the ERC Starting Grant COSI (grant no. 949808), a postdoctoral fellowship from the Special Research Fund (BOF) of Ghent University (BOF23/PDO/077 or BOF.PDO.2024.0008.01) and the Marie Skłodowska-Curie Actions (MSCA) postdoctoral fellowship INTERCOM (grant no. 101107007) to ESMF and IDC.

---

**Figure S1.**
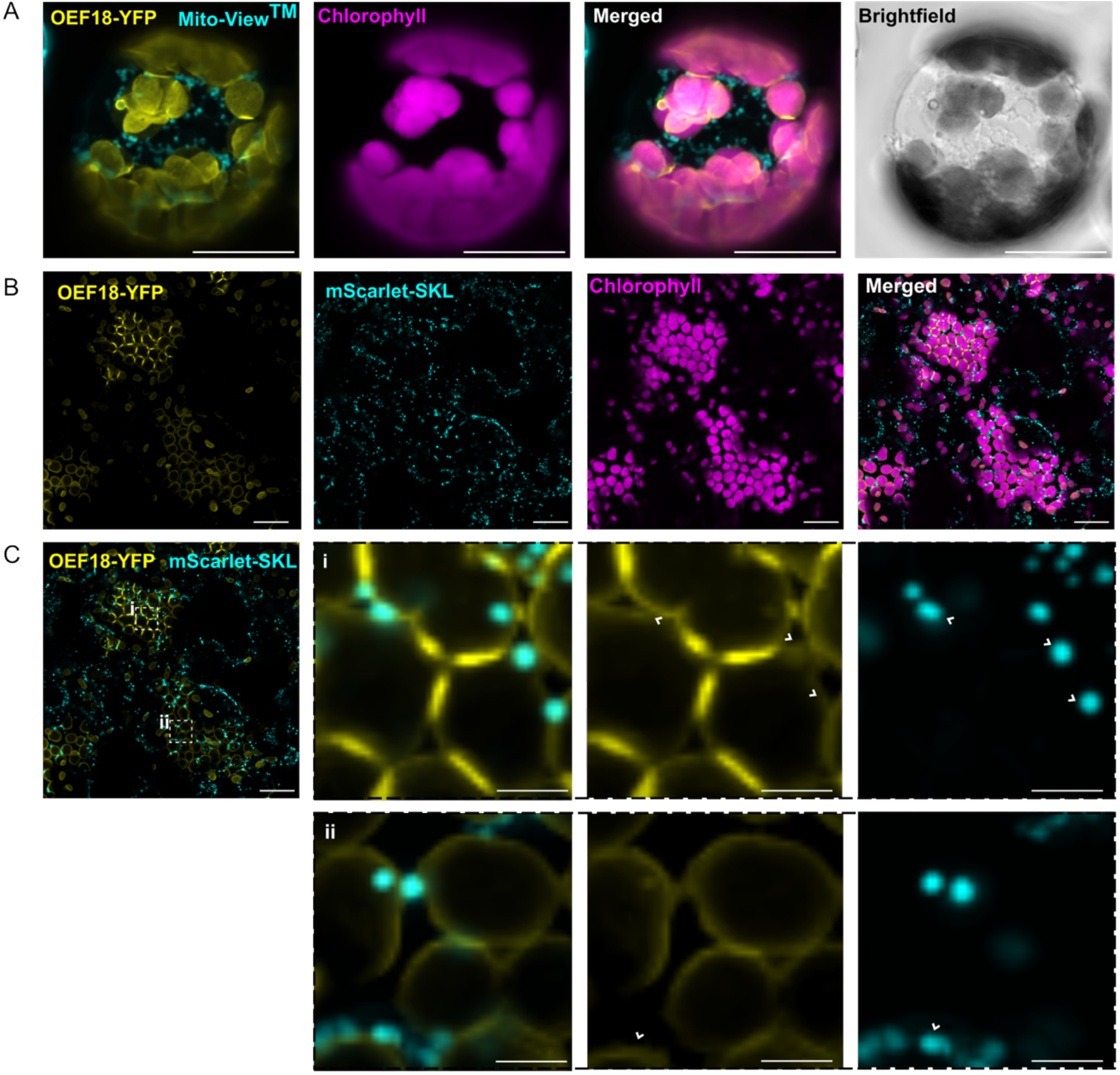
OEF18 localizes to the chloroplast rather than mitochondria or peroxisomes in *A. thaliana*. (A) Confocal microscopy image of an *A. thaliana* protoplast expressing OEF18-YFP. Mitochondria were stained with a MitoView™ 405 dye. Scale bar, 20 μm. (B, C) Co-localization of OEF18-YFP with the peroxisomal marker mScarlet-SKL in *A. thaliana* leaf epidermal cells. Scale bar, 20 μm. Insets (dashed boxes) show magnified views of selected regions labeled i and ii. Scale bar, 5 μm.

**Figure S2.**
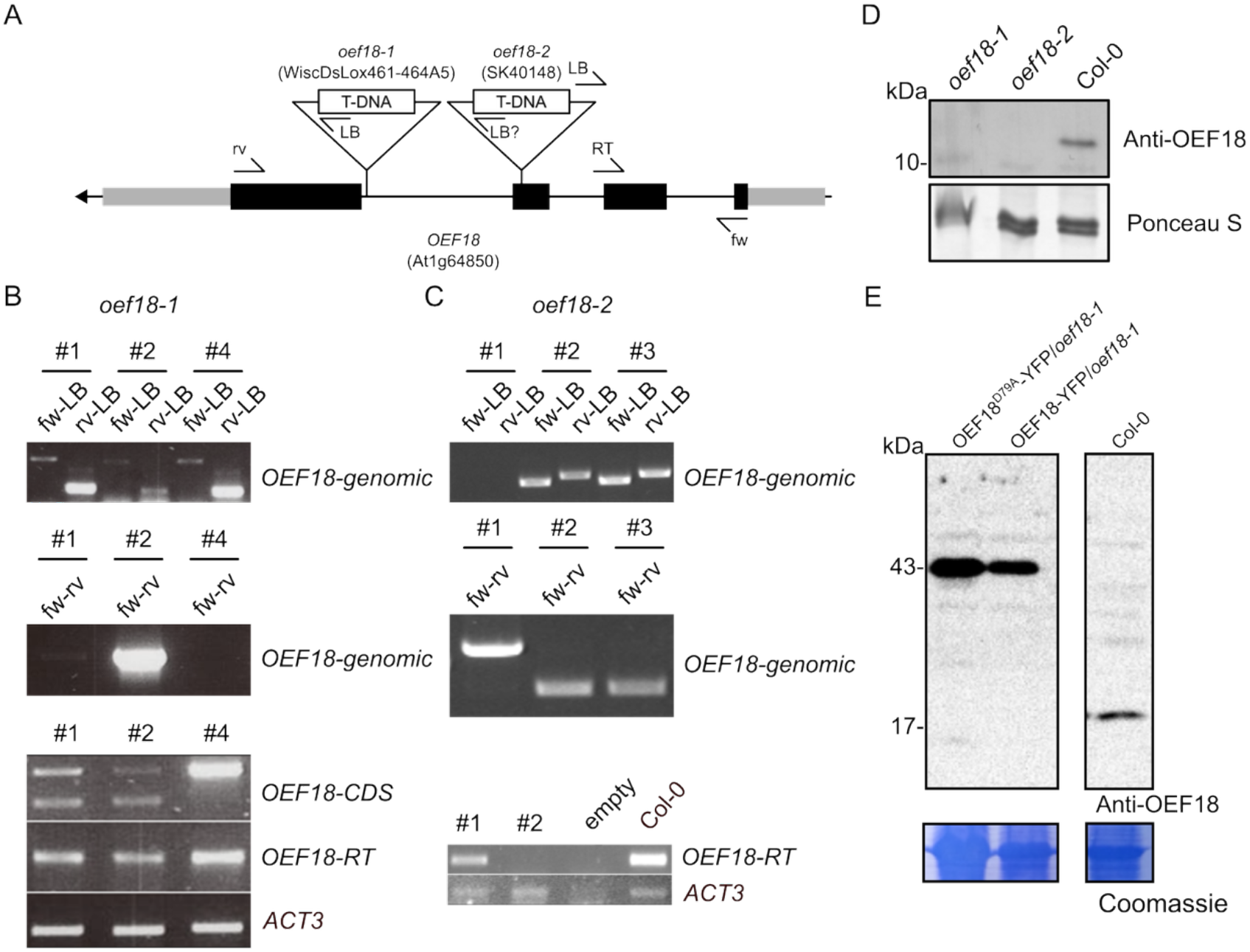
Genotyping of *oef18* T-DNA insertion alleles and OEF18 complementation lines. (A) Schematic representation of the *A. thaliana* OEF18 gene and the positions of T-DNA insertions. Exons are black. Primers used for the genotyping of the lines are indicated with fw (forward), rv (reverse) and RT (primers spans intron 2 for use with RT-PCR). (B) Genotyping of *oef18-1* by PCR. Top panel displays the search for positive T-DNA insertion lines on genomic DNA with fw and rv primers in combination with the left border primer of the T-DNA. 3 representative samples among various plants are shown (plant #1,2 and 4). Middle panel shows the search for homozygous T-DNA insertion lines. Bottom panel shows the product of RT-PCR for full-length OEF18 CDS (fw and rv primer) and N-terminal part of OEF18 (fw and RT primer). Actin3 (ACT3) is a loading control. (C) Genotyping of *oef18-2*, similarly to (B). In the bottom panel, the lane indicated with empty is a water control without cDNA. (D) Western blot against OEF18 in the *oef18-1* and *oef18-2* mutants and wild-type (Col-0). (E) Western blot against OEF18 in wild-type (Col-0) and OEF18 complementation lines in *A. thaliana* plants.

**Figure S3.**
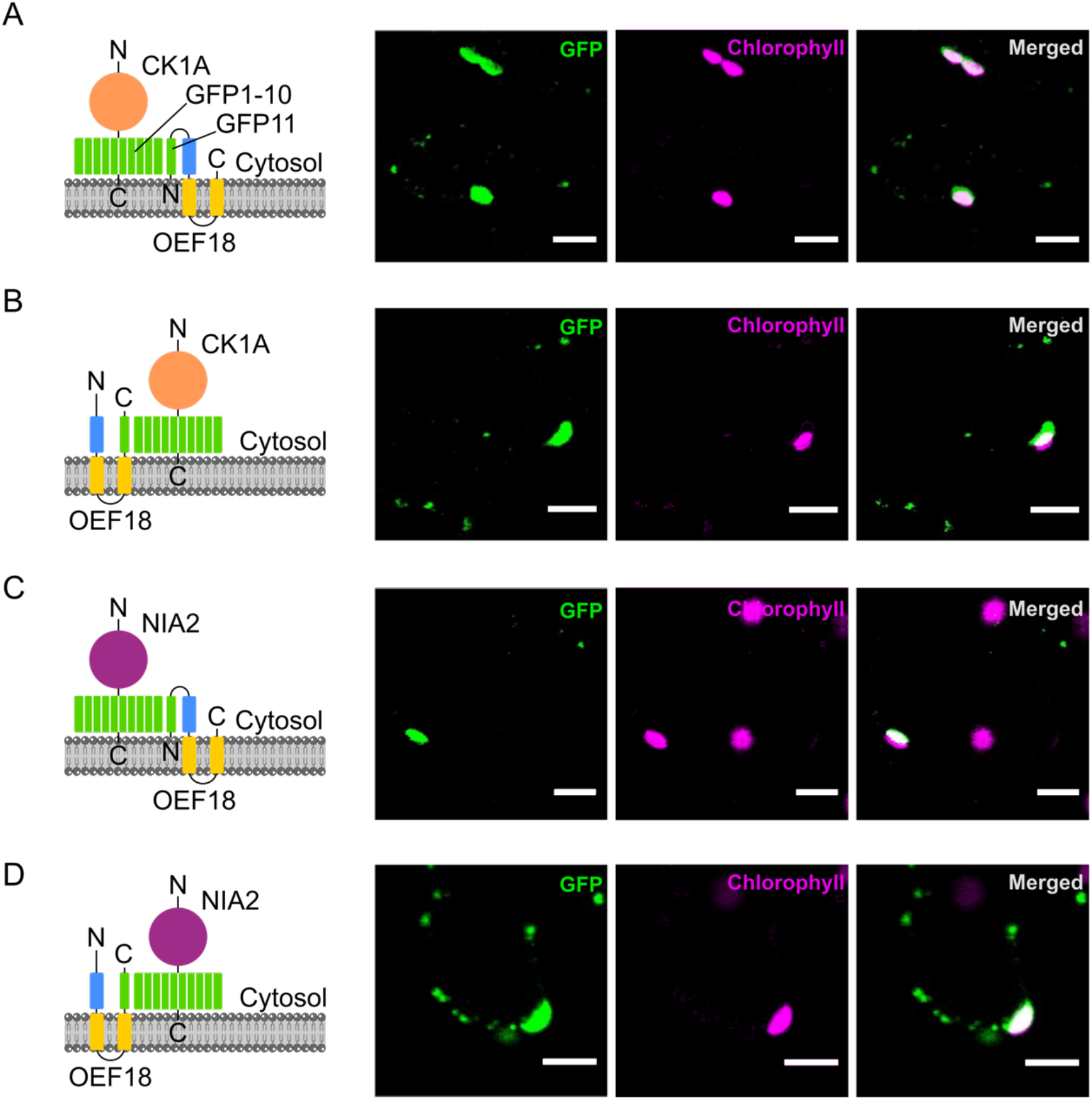
Both the N-terminus and C-terminus of OEF18 are localized in the cytosol. (A-D) Topology assessment of OEF18 with saGFP. Schematic diagrams of OEF18 constructs (left) and confocal microscopy images showing their subcellular localization (right). *N. tabacum* cells co-expressing either the cytosolic proteins CK1A_1-10C_ (A and B) or NIA2_1-10C_ (C and D) together with OEF18_11N_ (A and C) or OEF18_11C_ (B and D). Scale bar, 20 μm.

**Figure S4.**
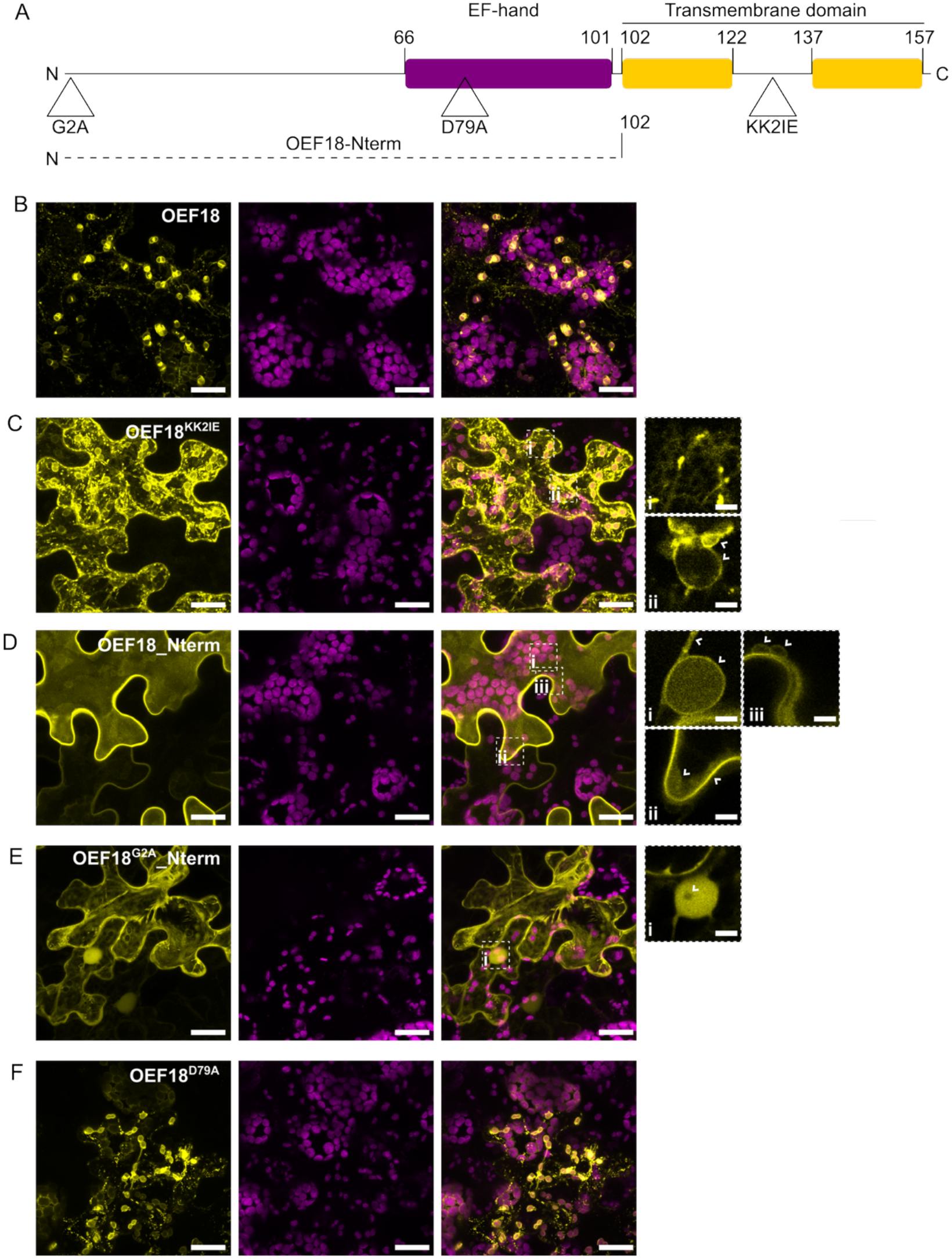
The Lys-Lys motif and N-terminal Gly are essential determinants of OEF18 localization. (A) Schematic representation of OEF18 and the mutated variants used in this study. The position of the N-terminus, EF-hand domain and the transmembrane domain are indicated. (B-F) Z-projections of confocal microscopy images of OEF18 and its variants fused to YFP in transient expression assays in *N. tabacum*. (B) OEF18-YFP co-localizes largely with chloroplast envelope, stromules, and slight punctate structures of which some likely are peroxisomes. (C) OEF18^KK2IE^-YFP, in which the Lys-Lys motif was mutated to Ile-Glu, still localized to the chloroplast envelope but additionally displayed prominent punctate accumulations of unknown origin and a more reticular distribution pattern suggestive of endoplasmic reticulum (ER) accumulation. This reticulate YFP signal is more clearly visible in single confocal optical sections shown in inset i, whereas inset ii highlights an ER-associated perinuclear accumulation pattern. (D) OEF18_Nterm-YFP, in which the transmembrane domain was deleted, exhibited a largely diffuse cytosolic distribution but retained evidence of membrane association. Representative examples are highlighted in the insets: i, perinuclear membrane association; ii, predominantly cytosolic accumulation with enhanced signal at or near the plasma membrane; and iii, residual association with the chloroplast envelope. (E) OEF18^G2A^_Nterm-YFP, which is identical to the construct shown in (D) but carries an additional Gly-to-Ala substitution at the N terminus, abolishing the predicted N-myristoylation site, exhibited an exclusively nucleo-cytoplasmic distribution. This was evidenced by diffuse cytoplasmic fluorescence together with clear nuclear accumulation, with the nucleolus indicated in inset i. (F) OEF18^D79A^-YFP, in which the Ca²⁺-coordinating Asp79 residue within the EF-hand motif was substituted to Ala, exhibited a subcellular localization pattern comparable to that of the wild-type OEF18-YFP construct. Scale bar, 20 μm.

**Figure S5.**
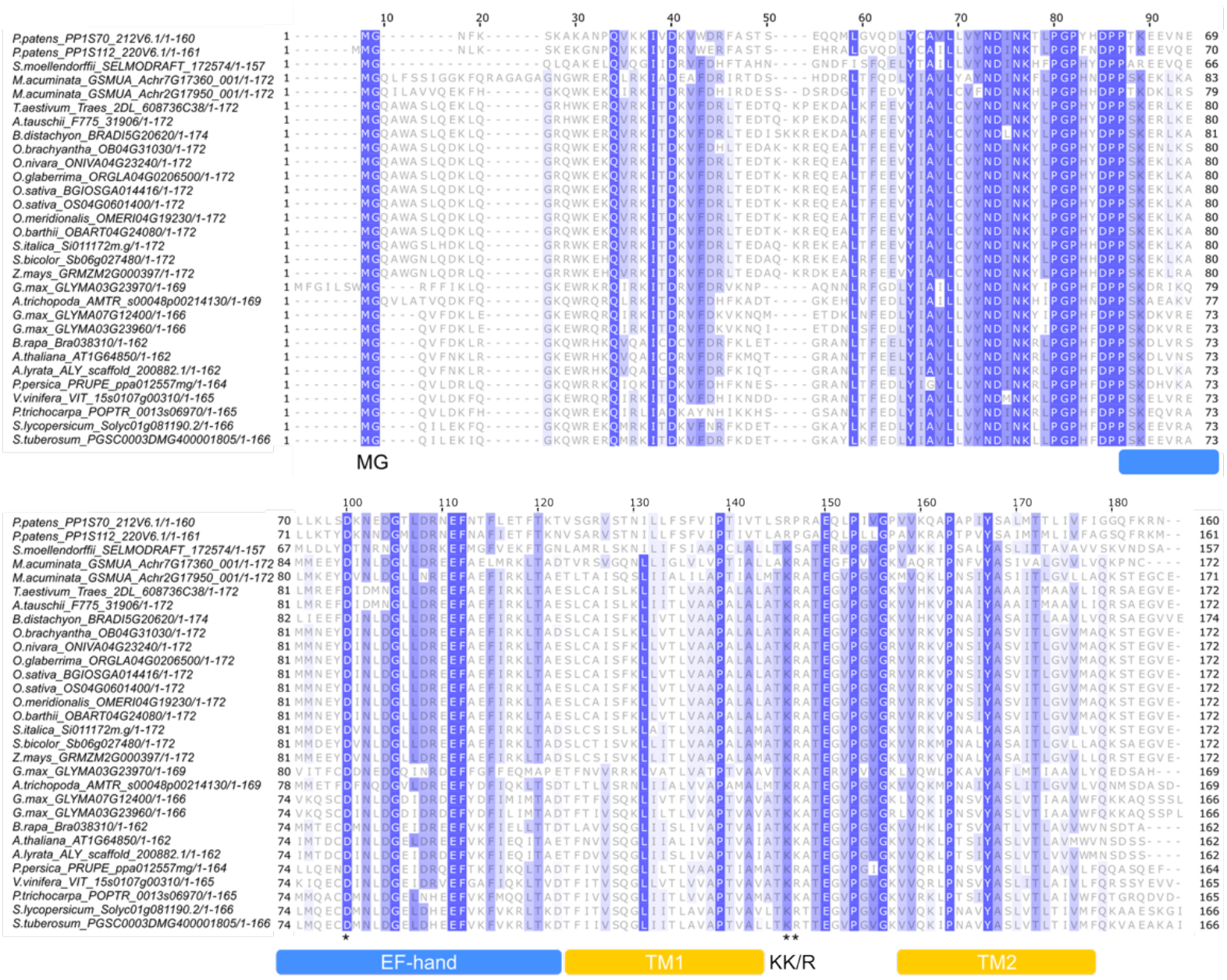
OEF18 and its orthologs exhibit conserved sequence domains. Alignment of OEF18 homologs from representative plant species. Amino acid shading intensity indicates the percentage of sequence identity. Sequence features are annotated based on *A. thaliana* OEF18, including an N-terminal MG motif, an EF-hand calcium-binding domain (predicted by PROSITE), two transmembrane helices (TM1 and TM2, predicted by ARAMEMNON), and a C-terminal KK/R motif.

**Figure S6.**
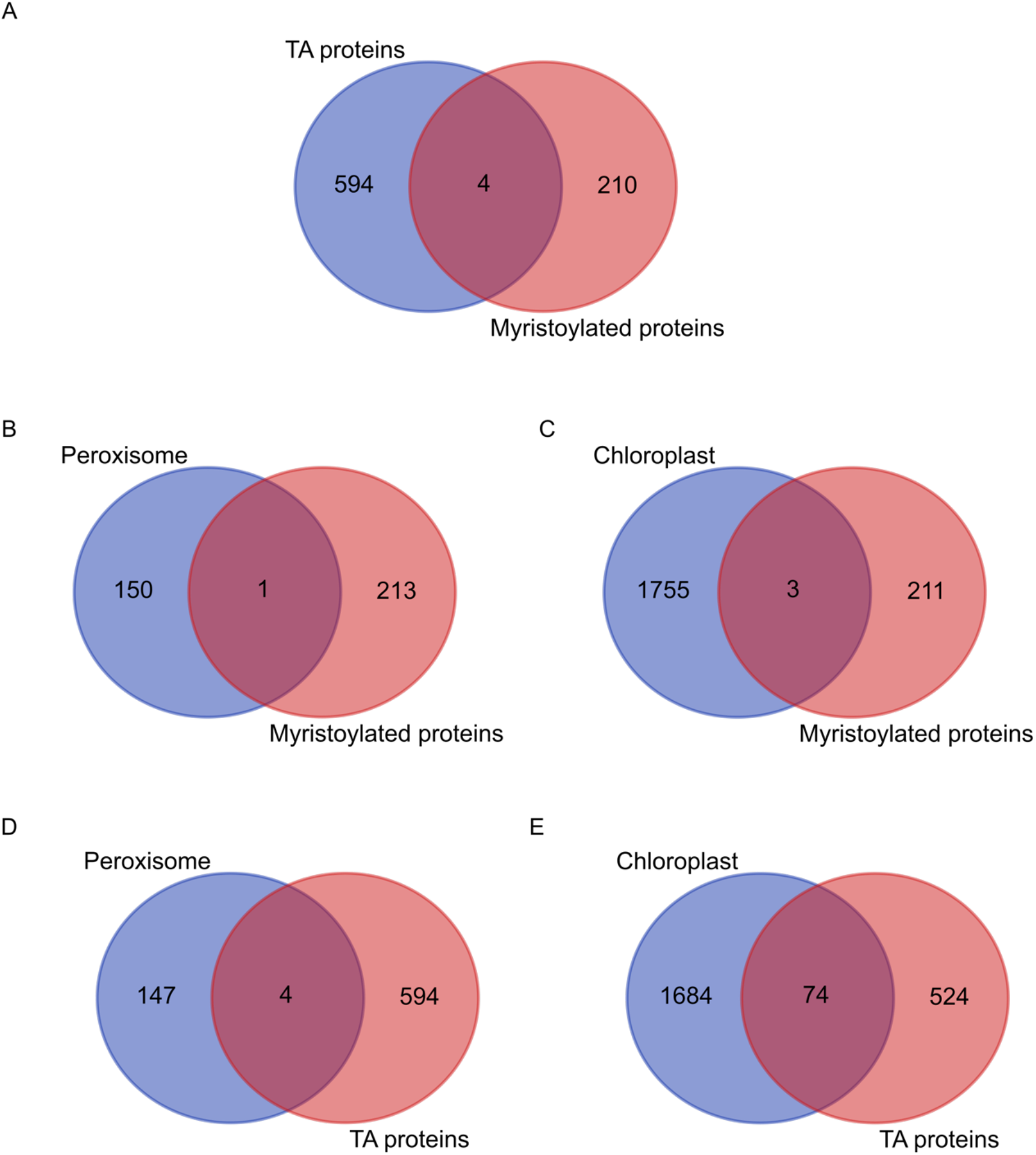
Venn diagram showing the overlap between myristoylated proteins, tail-anchored proteins and specific organelle-localized proteins. (A) A Venn diagram showing the overlap of identified proteins between TA proteins and myristoylated proteins. (B, C) Venn diagrams showing the overlap of identified proteins between myristoylated proteins and peroxisome-localized (B), chloroplast-localized (C) proteins. (D, E) Venn diagrams showing the overlap of identified proteins between TA proteins and peroxisome-localized (D), chloroplast-localized (E) proteins.

**Figure S7.**
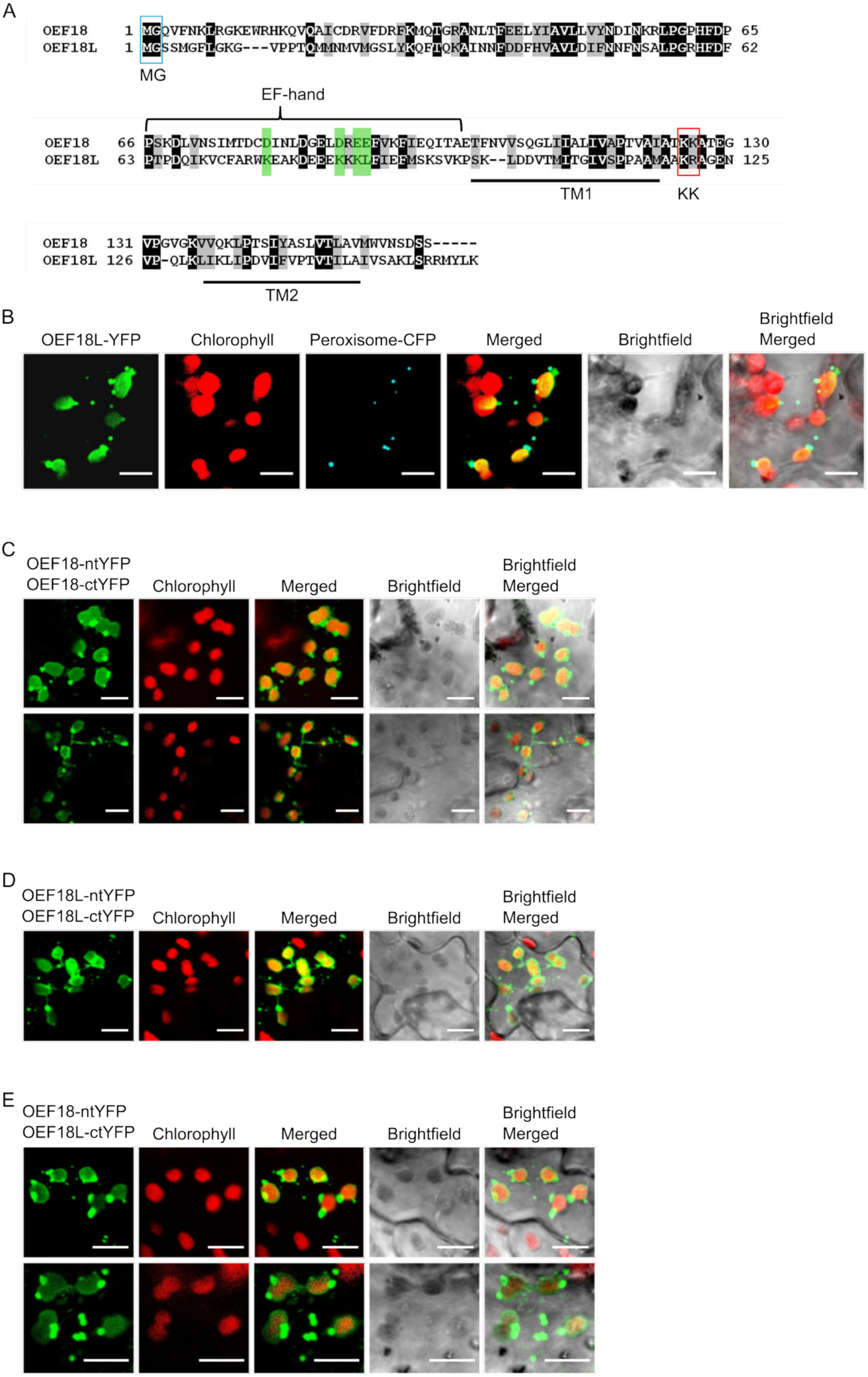
Similar to OEF18, OEF18L has conserved functional domains and localizes to the chloroplast envelope and co-localizes with a peroxisome marker in *N. tabacum*. (A) Sequence alignment of OEF18 and OEF18L proteins. Conserved amino acid residues are highlighted in black and similar amino acids in grey. Key structural features are indicated, including a conserved N-terminal MG motif (boxed in blue), an EF-hand calcium-binding domain (green highlights indicate critical binding residues in OEF18), two predicted transmembrane domains (TM1 and TM2, underlined), and a conserved C-terminal lysine pair motif (KK, boxed in red). (B) *N. tabacum* cells co-expressing OEF18L-YFP and peroxisome-CFP. Scale bar, 20 μm. (C-E) Transient BiFC assay in *N. tabacum* cells co-expressing OEF18-ntYFP and OEF18-ctYFP (C), OEF18L-ntYFP and OEF18L-ctYFP (D) or OEF18-ntYFP and OEF18L-ctYFP (E). Scale bar, 20 μm.

**Figure S8.**
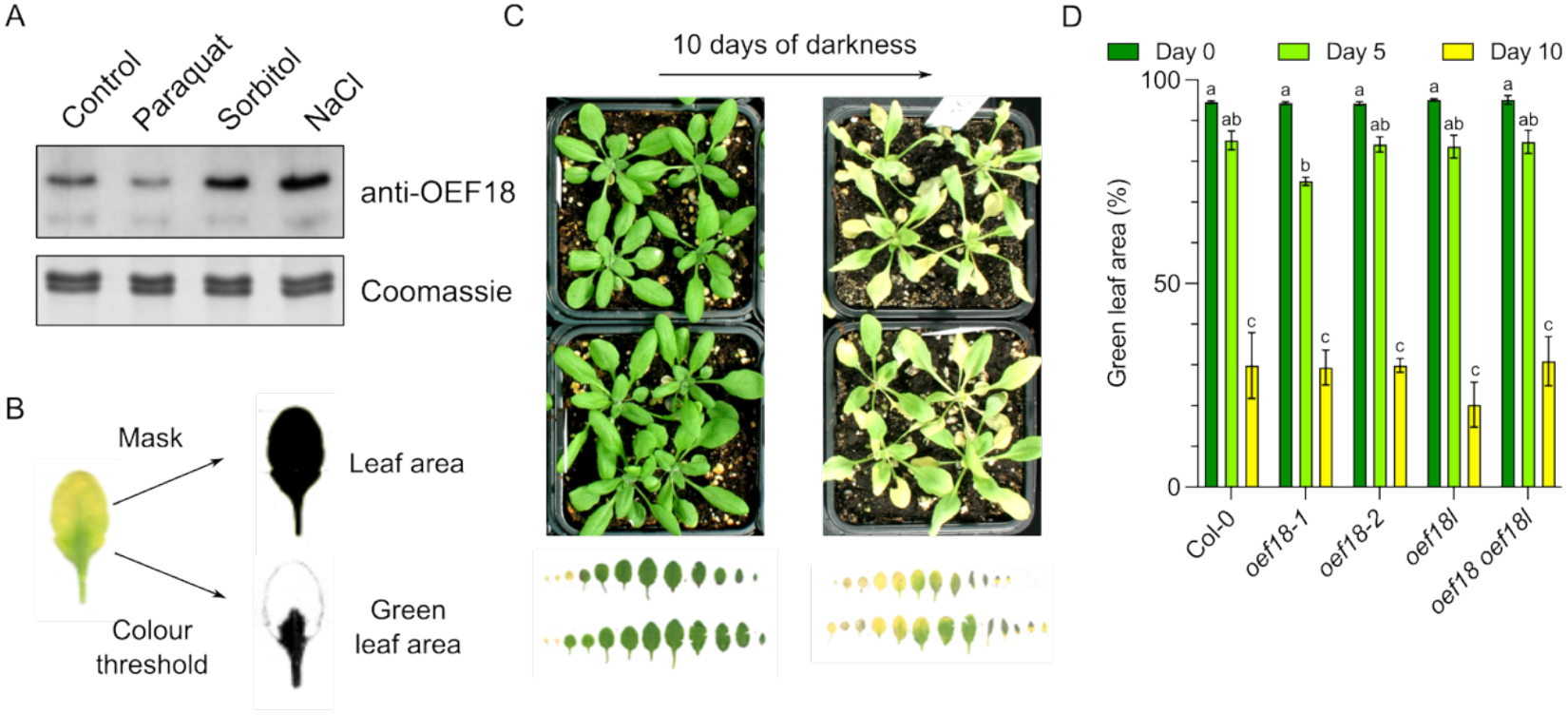
OEF18 is not involved in darkness-induced senescence. (A) Expression of OEF18 in complete seedlings (2 weeks) on agarose plates (1/2 MS, 1% sugar) under different stresses: reactive oxygen species (1μl/L paraquat), osmotic (100 mM sorbitol) and salt (75 mM NaCl). (B) The workflow of image analysis in ImageJ software to calculate the green leaf area. (C) Top view of *Arabidopsis* plants grown in soil for 4 weeks under long-day conditions and then transferred to darkness for 10 days. (D) Quantification of green leaf area (%) of the indicated genotypes after darkness treatment. Data represent mean ± SD. Different letters indicate statistically significant differences (P < 0.05, two-way ANOVA with Tukey’s test).

**Figure S9.**
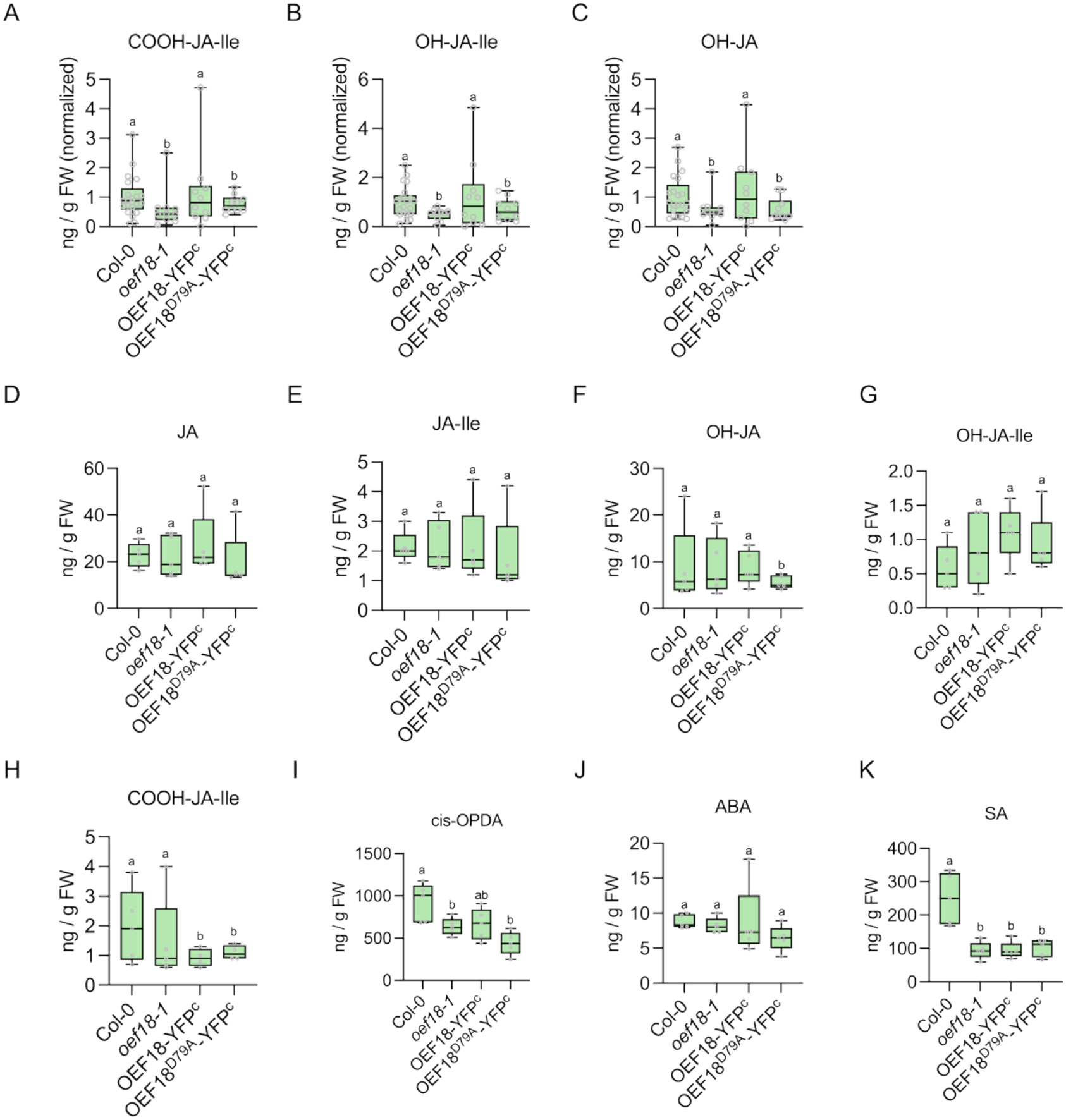
Hormone and related metabolite levels in *oef18-1* and complemented genotypes. (A-C) Concentration of COO-JA-lle (A), OH-JA-lle (B), OH-JA (C) in Col-0 compared to *oef18-1*, OEF18-YFP^C^, OEF18^D79A^-YFP^C^ after three hours of feeding by fourth instar *S. littoralis* larvae. (D-K) Concentration of JA (D), JA-lle (E), OH-JA (F), OH-JA-lle (G), COO-JA-lle (H), cis-OPDA (I), ABA (J) and SA (K) in Col-0 compared to *oef18-1*, OEF18-YFP^C^, OEF18^D79A^-YFP^C^ during steady-state conditions of *A. thaliana* grown on soil. Different letters indicate statistically significant differences (P < 0.05, one-way ANOVA with Tukey’s test).

